# Comprehensive characterization of toxins defines progression and stages in a non-human primate model of inhalation anthrax

**DOI:** 10.1101/2022.07.13.499850

**Authors:** Anne E. Boyer, Maribel G. Candela, Renato C. Lins, Maria I. Solano, Adrian R. Woolfitt, John Lee, Daniel C. Sanford, Katherine Knostman, Conrad. P. Quinn, Alex R. Hoffmaster, James L. Pirkle, John R. Barr

## Abstract

Inhalation anthrax has three clinical stages: early-prodromal, intermediate-progressive and late- fulminant. The toxins produced during infection exert pathologic effects leading to death, but they have not been comprehensively characterized throughout the course of infection. Mass spectrometry methods for anthrax toxins, total-protective antigen (PA), total-lethal factor (LF), total-edema factor (EF), and toxin complexes, lethal toxin and edema toxin were used to characterize the stages of inhalation anthrax in 23 cynomolgus macaques. The target aerosol dose was 200 LD_50_ *B. anthracis* Ames spores. 22 animals died during the study. Different patterns of toxemia and bacteremia were observed in 11 animals with the shortest survival times (fast progression), the 11 animals with longer survival (slow progression), and the one survivor. Toxemia and bacteremia were predominantly triphasic with an early rise (phase-1), a plateau/decline (phase-2), and a final rapid rise (phase-3). The patterns were consistent for all toxins. The end-of-phase-1 LF was higher in fast progression [median(lower quartile– upper quartile)] of [195(57.4–326)-ng/mL], than in slow progression [23.8(15.6–26.3)-ng/mL] (p=0.0001), or the surviving animal [11.1-ng/mL]. End-of-phase-1 EF was also higher in fast [22.2(2.7– 42.8)-ng/mL] than slow progression [0.17(0.064–0.066)-ng/mL] (p=0.0005), or the surviving animal [0.040-ng/mL]. Animals with slow progression and lower end-of-phase-1 toxemia, had an extended plateau/decline (≥24-hours), with low variability of PA, LF, and LTx across all animals. Its characterization revealed an upper threshold; a limit for exiting phase-2 and entering the critical phase-3, 342-ng/mL (PA), 35.8-ng/mL (LF), and 1.10-ng/mL (EF). The thresholds were exceeded early in animals with fast progression (38.3±7.4-hours) and later in slow progression (78.7±14.1-hours). Once the threshold was passed toxin levels rose rapidly, differences in toxemia were reduced, and the duration to terminal was rapid and similar; 21.0±7.3-hours for fast and 20.4±7.3-hours for slow. This first comprehensive evaluation of anthrax toxins defined all stages, providing insights into disease progression.

**Author Summary:** The comprehensive analysis of all major anthrax toxins and bacteremia in a non-human primate model of inhalation anthrax revealed a triphasic kinetics of toxemia that aligns with the three clinical stages, early-prodromal, intermediate-progressive and late-fulminant. End of phase-1 toxin levels may predict the subsequent speed of progression. Phase-2 toxemia helped define critical thresholds representing the entry to phase-3. Exceeding these thresholds was associated with a short remaining survival time (about 21 hours). This first comprehensive characterization of toxemia provides knowledge and guidance for better management of anthrax.

## Introduction

*Bacillus anthracis* is a gram positive, spore-forming bacterium and the causative agent of anthrax (1). Anthrax can take many forms, cutaneous, gastrointestinal, injection, and the most lethal, inhalation. As such, *B. anthracis* is a U.S. Category A priority pathogen, and Tier 1 select agent (www.selectagents.gov), with the potential for aerosol dispersion and mass casualties (2). Historically, the fatality rate of untreated inhalation anthrax ranged from 89%-100% (3–6). In 2001, anthrax spores were intentionally distributed through the U.S. mail. Even with the use of antimicrobial agents and modern intensive medical support, the fatality rate was 45% (7–9). The high rate of fatality is due to the production of two binary toxins that disable host’s immune response, cause endothelial barrier dysfunction, edema, and hemorrhage (10). Even after the clearance of bacilli by antimicrobial treatment, the toxins persists for days, causing further damage and death (11). A better understanding of the levels of anthrax toxins during the stages of infection provides insights into their role in pathogenicity.

The two major *B. anthracis* toxins consist of binary combinations of protective antigen (PA), which binds enzymatic factors, lethal factor (LF) and edema factor (EF), forming active toxin complexes lethal toxin (LTx) and edema toxin (ETx), respectively (12). PA is secreted as an 83-kDa protein (PA83) that binds cell surface receptors TEM8 and CMG2 (13, 14). There it is hydrolyzed by furin-like proteases to 63 kDa (PA63), which causes it to form heptamers/octamers that readily bind LF and EF, forming the LTx and ETx complexes (15–18). PA83 is also hydrolyzed by serum proteases, forming PA63, and ultimately, LTx and ETx (19–23). LF, a 90-kDa zinc-dependent endoproteinase, hydrolyzes and inactivates several members of the mitogen-activated protein kinase kinase (MAPKK) family of proteins that are central to various cell signaling responses (24). EF, an 89-kDa calcium and calmodulin dependent adenylate cyclase, has a high rate of catalysis of ATP to cyclic AMP (25). The accumulation of cyclic AMP contributes locally to edema, but also activates protein kinase A and exchange protein, both of which control a variety of cellular functions. In early stages of infection, LTx and ETx synergistically suppress innate immune responses, allowing the infection to take hold, bacteria to proliferate, and toxins to accumulate (10, 26–28). In later stages the toxins contribute to disruption of endothelial barrier integrity, causing edema, pleural effusions, hemorrhage, and hemodynamic instability (29–31). After a certain point, the damage becomes insurmountable and largely resistant to standard procedures of cardiovascular support (32). It is not yet known what toxin levels correlate with these stages and their clinical impact.

Clinically, inhalation anthrax has been described as having three-stages: 1) the early-prodromal stage, 2) intermediate-progressive stage, and 3) late-fulminant stage (33, 34). The first stage presents with non-specific symptoms, often resembling the flu. The cure rate is highest in this early stage, but since anthrax is rare, a low index of suspicion at clinical presentation can delay appropriate treatment. In the second stage, non-specific symptoms continue but pathology and the clinical signs progress. The potential for survival is still high in this stage but requires aggressive medical intervention. Onset of the third stage is rapid with respiratory failure, shock, and often meningitis. The likelihood of survival is low in the third stage. This was most evident during the anthrax letter attacks of 2001. All 11 cases with inhalation anthrax received aggressive medical intervention. However, it was the stage at presentation that was associated with outcome. The six that presented during stage-2 survived, while the five presenting in stage-3 did not (7, 8, 34). Novel treatments such as antitoxins developed in more recent years may have helped to improve outcome (35). However, more needs to be learned about the progression of anthrax to help discern stages.

Of the animal models for inhalation anthrax, non-human primates (NHP), both rhesus macaques and cynomolgus macaques have been extensively studied for anthrax pathology, therapeutics, and vaccines (36–43). However, of the many toxins and biomarkers of anthrax, only PA and bacteremia have been characterized in cynomolgus macaques (42). We have developed and validated sensitive and specific mass spectrometry (MS) methods for all major anthrax toxin proteins and their toxin complexes. They include methods for total-LF (LF+LTx) (44), LTx (PA-LF) (22), total-EF (EF+ETx), ETx (PA-EF) (23), and PA (total PA, full length PA83, and PA63) (45). Application of the methods in five rhesus macaques showed a triphasic kinetics of toxemia (22, 23, 45, 46). These were too few and limited in scope to draw major conclusions about staging. Here, these MS methods were applied to measure and characterize anthrax toxins over the course of infection in the cynomolgus macaque NHP model. This study provides the first comprehensive assessment of all the major anthrax toxins during infection which, along with traditionally measured bacteremia, addresses the complexities of anthrax progression, yielding insights that may improve medical management and public health outcomes during an exposure event.

## Results

### 1. Cynomolgus macaque study outcomes

#### Exposures and clinical outcomes

The study included 23 cynomolgus macaques, 12 males and 11 females, with a mean±standard deviation weight of 4.0±0.61 kg at challenge (Table 1). The weight for females was 3.7±0.48 kg and males was 4.3±0.61 kg. The NHP’s were randomized and challenged in two groups. Group-1 included 10 animals with two non-challenged control, for 8 challenged in group-1. The two controls did not develop disease or detectable toxemia/bacteremia. Group-2 included the two control animals from group-1 which were challenged with 13 others, for 15 animals. The collective exposure dose of 221(±29.2) Ames LD_50_ and that of both group-1 (218±32 LD_50_) and group-2 (223±29 LD_50_) were similar. There were 22 non-survivors, ordered by time to death or euthanasia which ranged from 48.5-201.9 hours-h (Table 1 and S1 Fig, survival curve). One animal survived to the end-of-study.

**Table 1.** Cynomolgus macaques, challenge, symptoms, and survival time.

|  | Animal ID | Gender | Weight (Kg) Day 0 | Ames LD<sub>50</sub> | LD<sub>50</sub> per kg | Symptom Onset-h | First Symptom | Hours to Death (Days) | Manner of Death | Pathology Cause of Death |
| --- | --- | --- | --- | --- | --- | --- | --- | --- | --- | --- |
| 1 | C58282<sup>1</sup> | M | 4.6 | 219 | **48** | <u>29.1</u> | HP | 48.5 (2.0) | Euth | Sepsis |
| 2 | C59621 | F | 3.2 | 230 | 72 | 47.4 | N/Eu | 48.9 (2.0) | FD | Sepsis |
| 3 | C60763 | M | 3.3 | 200 | 61 | 54.7 | FD | 54.7 (2.3) | FD | Sepsis |
| 4 | C58176 | M | 4.2 | 280 | 67 | 38.2 | W | 55.2 (2.3) | FD | Sepsis |
| 5 | C58003<sup>+</sup> | F | 4.2 | 232 | 55 | 40.3 | HP | 57.3 (2.4) | FD | Sepsis |
| 6 | C59656 | F | 3.7 | 206 | 56 | 40.9 | HP | 57.9 (2.4) | FD | Sepsis |
| 7 | C58214<sup>1</sup> | F | 3.4 | 246 | 72 | <u>28.5</u> | HP | 58.4 (2.4) | FD | Sepsis |
| 8 | C60732 | M | 3.1 | 258 | 83 | 35.8 | HP | 59.0 (2.5) | Euth | Sepsis |
| 9 | C59669 | F | 3.4 | 223 | 66 | 56.3 | HP, L | 64.8 (2.7) | Euth | Sepsis |
| 10 | C57161<sup>1</sup> | F | 3.7 | 203 | 55 | 41.5 | HP | 66.9 (2.8) | FD | Sepsis |
| 11 | C58170<sup>+</sup> | M | 4.8 | 208 | 43 | 61.4 | HP | 74.1 (3.1) | Euth | Sepsis |
| 12 | C59619 | F | 3.6 | 270 | 75 | 35.3 | W, HP, V | 76.4 (3.2) | FD | Sepsis |
| 13 | C58135 | M | 4.0 | 186 | 47 | <u>34.9</u> | HP | 90.4 (3.8) | Euth | Sepsis |
| 14 | C57793 | M | 4.1 | 187 | 46 | 67.0 | FA | 91.1 (3.8) | FD | Sepsis |
| 15 | C58167 | M | 4.9 | 215 | 44 | 37.3 | HP, W | 91.7 (3.8) | FD | Sepsis |
| 16 | C57998<sup>1</sup> | F | 3.5 | 199 | 57 | 77.2 | L, W | 100.8 (4.2) | FD | Sepsis |
| 17 | C59617 | F | 4.9 | 203 | 41 | 88.1 | HP | 105.1 (4.4) | FD | Meningitis / Sepsis |
| 18 | C58159<sup>1</sup> | M | 4.2 | 188 | 45 | <u>30.2</u> | HP | 109.0 (4.5) | Euth | Meningitis |
| 19 | C58122<sup>1</sup> | M | 4.8 | 181 | 38 | 35.6 | DI | 115.1 (4.8) | FD | Sepsis |
| 20 | C57997<sup>1</sup> | F | 3.9 | 234 | 60 | 53.9 | W | 125.9 (5.2) | FD | Sepsis |
| 21 | C58156<sup>1</sup> | M | 4.7 | 274 | 58 | 60.7 | HP | 140.1 (5.8) | FD | Meningitis |
| 22 | C59618 | F | 3.4 | 202 | 60 | 57.0 | HP | 201.9 (8.4) | Euth | Sepsis |
| 23 | C58174 | M | 4.9 | 239 | **49** | 108.4 | FA | 336.6 (14) | EOS | NA |
|  | Mean±SD |  | 4.0±0.61 | 221±29.2 | 56.4±12.0 | 50.4±20.2 |  | 86.1±37.1 |  |  |

23 animals were challenged in two groups. <sup>1</sup>First group challenged included 8 challenged animals and 2 non-challenged controls. The second group challenged included 15 animals, among these were the 2 controls from the first group<sup>+</sup>. SD-standard deviation. LD<sub>50</sub>/Kg in bold font show similar exposure doses for the animal with shortest survival and the surviving CM. \*Survivor sacrificed at the end-of-study (EOS), 14 days. Animals were observed every 6 hours for symptoms and criteria for euthanasia. Symptom onset is the time from challenge to first symptom. Symptoms included hunched posture (HP), wheezing (W), lethargic (L), vomiting (V), facial edema (FA) and diarrhea (DI), or none (N) until euthanized (E), found dead (FD).

When adjusted by weight, the Ames LD_50_ exposure dose (LD_50_/kg) tended to be higher in animals that had shorter survival times with some notable exceptions. The animal with the shortest survival time (48.5 hours) dose (48 LD_50_/kg) was similar to the animal that survived (49 LD_50_/kg), and lower than the animal with the latest time to death (60 LD_50_/kg). The mean±standard deviation spore dose/kg of the 11 animals with shorter survival times was 61.6±11.7 LD_50_/kg and for the 11 non-survivors with longer survival was 51.9±11.0 LD_50_/kg. These differences did not meet the alpha level of 0.05 for significance (p=0.0583). As can be seen in Table 1, a longer time to death does not correlate with lower doses.

#### Symptom onset

The onset of symptoms has been represented by an increase in body temperature from normal or baseline (47). Body temperatures were monitored by telemetry before and after challenge. However, the increase in body temperature was not a consistent indicator of symptoms. It was present in only 12 animals for the shoulder transponder and 11 animals for the hip. When present, the elevated temperatures were brief, even transient in most, except in the shoulder of three animals with the longest survival time (S2 Fig).

Symptoms were also monitored visually by veterinarians, every 6 hours after challenge. The first clinical sign(s) and the time of occurrence (symptom onset) was recorded (Table 1). The time from challenge to symptom onset ranged from 28.5 hours at the earliest, to 108.4 hours at the latest, in the survivor. Mean±standard deviation time to symptom onset was 50.4±20.2 hours. The most common first clinical sign was hunched posture, present in 15 animals and the least common was diarrhea and vomiting, present in only one each. Four animals had two or more symptoms at onset, while two had no clinical signs recorded before early death/euthanasia at 48.9- and 54.7-hours, suggesting the progression from normal behavior to death or moribund occurred in less than 6 hours.

#### Cause of death

Terminal microscopic and gross pathology, to be described later, provided the final determination of the cause of death as being due to either sepsis, for which bacteremia was found in multiple organs, or meningitis, for which whole brain pathology was primary and bacteremia was limited to lymphatic organs (Table 1). Only three animals, all with longer survival times, showed whole brain pathology; two of these were considered meningitis alone (pathology details below).

### 2. Anthrax toxemia and bacteremia

#### 2.1 Pre-symptom onset detection of anthrax biomarkers

The limits of detection for all the anthrax toxin methods are summarized in Table 2. Total-LF and -EF methods detect and measure the enzymatic activity which gives lower detection limits, whereas the PA method detects and measures the specific trypsin-digest peptide fragments of PA itself which has a higher detection limit. Figure 1 compares earliest detection of total toxins LF, EF, and PA, and bacteremia relative to symptom onset. This showed that LF was the first biomarker detected and it preceded symptom onset in all exposed animals. EF was the next biomarker detected and was first observed at the same time as LF in 9 of the animals, after LF but still before symptoms in 13 animals, and at symptom onset in one animal. PA and bacteremia were detected later with bacteremia detection preceding PA detection in 12 animals and coinciding in 11 animals. Bacteremia and PA detection was before symptom onset in 18 and 16 animals, respectively.

**Figure 1.**
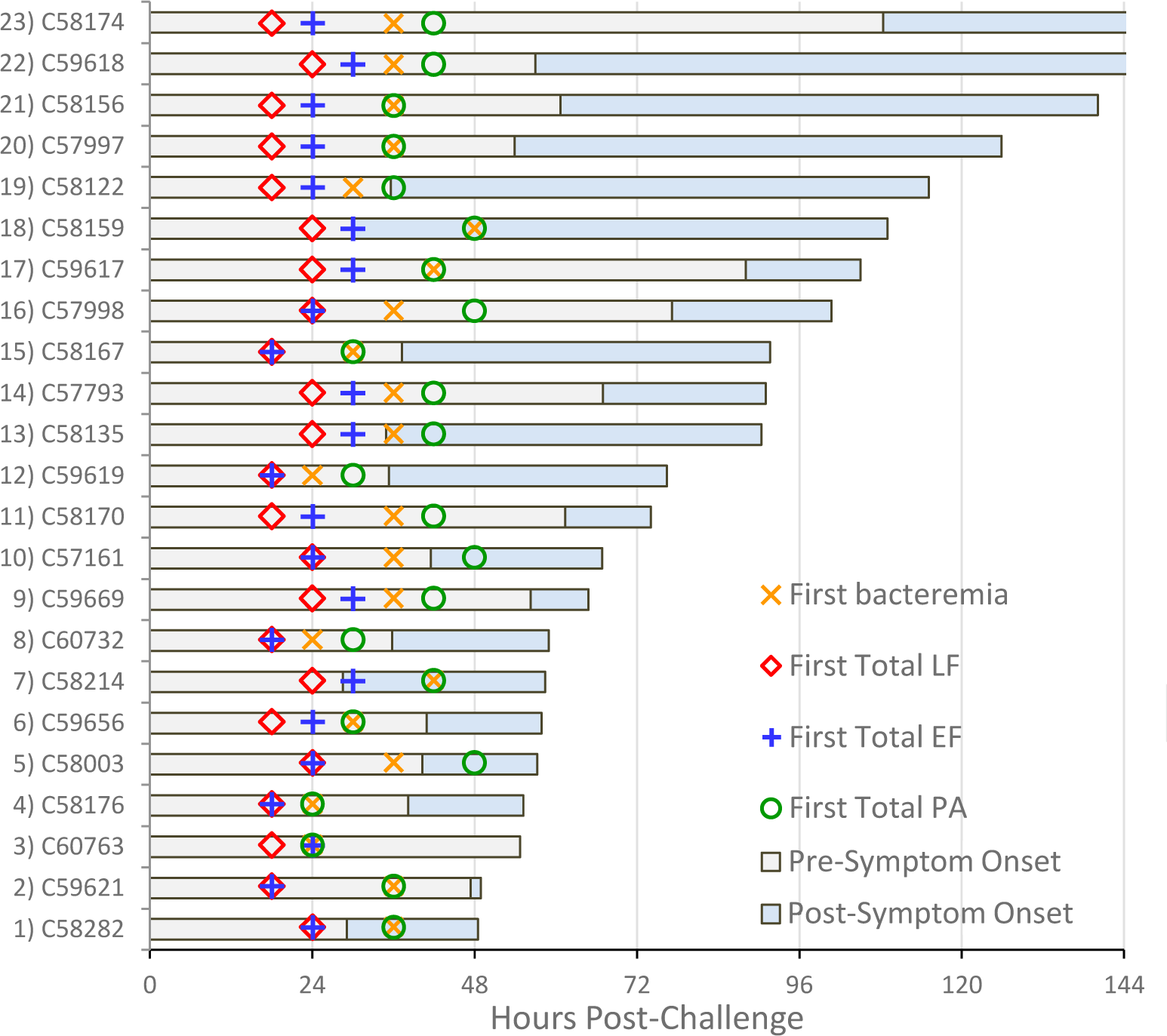
First detection of anthrax biomarkers. First detection of biomarkers relative to symptom onset in 23 cynomolgus macaques with inhalation anthrax. Pre-symptom onset period (gray bar) and post-symptom onset period (blue bar) out to 144 hours, for total lethal factor (LF), total edema factor (EF), total protective antigen (PA), and confirmed bacteremia. Detection limits for LF (0.0027 ng/mL), EF (0.00002 ng/mL), PA (1.87 ng/mL) and confirmed bacteremia (reported to 25 cfu/mL). Non-confirmed results of ‘+’ excluded here but included in Table 2.

**Table 2.** Anthrax toxin mass spectrometry (MS) 3-step methods summary.

| Analyte | Pre-MS<br>Sample Prep | LOD<br>(ng/mL) | pmol/mL |
| --- | --- | --- | --- |
| LF Total | IP-catalysis | 0.0027 | NA |
| LTx | IP-catalysis | 0.0075 | NA |
| EF Total | IP-catalysis | 0.00002 | NA |
| ETx | IP-catalysis | 0.00002 | NA |
| PA Total | IP-tryp digest | 1.87 | 0.0225 |
| PA83 | IP-tryp digest | 1.22 | 0.0147 |

Immunoprecipitation (IP), limit of detection (LOD), tryptic (tryp).

Table 3 includes the summary for pre-symptom onset detection for total toxins, toxin complexes, LTx and ETx, and full length PA83 for 1) the mean time to first detection, 2) the time that first detection occurred before symptom onset, and 3) the percent of animals positive for each biomarker at each early time point. It showed that LF was detected earliest, well before symptoms, and most frequently, in 100% of animals by 24 hours. EF was detected second earliest and most frequently, in 65% by 24 hours and 100% by 30 hours. Detection of bacteremia followed. Toxin complexes, LTx and ETx, were detected on average about 9 hours after their total-LF and total-EF counterparts and may reflect the accumulation and activation of PA83 to PA63, binding LF and EF in circulation. Total PA was detected later, even though PA was present earlier in the form of LTx and ETx. Detection limits are lower for the two enzymatic methods, allowing detection of lower levels earlier. Full length PA83 was the last to test positive and never reached 100% positivity at any time point for available samples. Later results show that PA83, when detectable, was only a small fraction of the total PA measured. Therefore, most of the PA was in the PA63 form throughout the entire course of infection.

**Table 3.** Pre-symptom onset detection.

| Biomarker | Time first detected (h) | Time detected before symptoms (h) | Percent positive at post-challenge time |  |  |  |  |
| --- | --- | --- | --- | --- | --- | --- | --- |
|  |  |  | 18h | 24h | 30h | 36h | 48h |
| Total LF | 20.9±3.1 | 29.6±20.5 | 52.2 | 100.0 | 100 | 100 | 100 |
| Total EF | 24.5±4.4 | 25.9±19.8 | 21.7 | 65.2 | 100 | 100 | 100 |
| Bact NC (+) | 30.0±7.0 | 20.4±19.1 | 17.4 | 26.1 | 52.2 | 100 | 100 |
| LTx | 30.5±4.4 | 19.9±19.5 | 0 | 22.7 | 100 | 100 | 100 |
| ETx | 33.4±7.6 | 17.0±22.6 | 0 | 21.7 | 52.2 | 73.9 | 100 |
| Bact Confirmed | 34.2±6.1 | 16.2±19.0 | 0 | 17.4 | 30.4 | 87.0 | 100 |
| Total PA | 38.1±7.4 | 12.3±19.7 | 0 | 8.70 | 26.1 | 50.0 | 100 |
| PA83 (n=21) | 59.9±39.1 | -11.2±36.2 | 0 | 0 | 4.3 | 18.2 | 40.9 |

First appearance of anthrax toxins and bacteremia. Mean time (hours-h) each biomarker was first detected post-challenge and the mean time (h) the first detected occurred before symptom onset in each animal. The percent positive for each biomarker of 23 animals at all early time points (prior to the mean symptom onset at 50.4 hours). Bacteremia was quantified by serial dilution, plating in triplicate and colony counting which has an LOD of 100 cfu/mL. Only bacteremia reported ≥100 cfu/mL were confirmed. Results below this were reported as + for two of three plates with less than 10 colonies.

Inclusion of the non-confirmed (NC) + results were included for Bact NC (+).

#### 2.2 Biomarker levels at symptom onset

Toxin levels measured at the time points closest to symptom onset were determined for each animal (Table 4). LF, LTx and EF were positive at symptom onset in all 23 animals (100%). Although the detection of LTx in all animals at symptom onset indicates that PA is present (as PA-LF), total PA, with a higher detection limit, was not detected in four animals in which the LTx levels were below 0.26 ng/mL. Total PA was detectable in 19 of the 23 animals (83%) at symptom onset, one of which had lower LTx of 0.893 ng/mL. ETx and bacteremia were detectable in 20 and 21 of 23 animals, respectively. PA83 was only positive in 7 animals (30.4%). Median symptom onset levels for all 23 animals were in the mid-low ng/mL for PA, LF, and LTx and sub-ng/mL for EF and ETx. The median for bacteremia was 1.3E+04 cfu/mL. Lower and upper quartiles for all 23 animals, demonstrate a broad range which was due to either or both the variable rates of progression and the different stages during which symptoms were first observed. As related to progression, median symptom onset levels were higher in 11 animals that died at earlier time points. The median symptom onset levels of PA, LF, LTx and bacteremia were 2.3-, 10-, 14-, and 22-fold higher and EF and ETx were 91- and 136-fold higher, respectively, for the 11 animals with the shortest survival times, than the 11 animals with longer survival times. As related to time of symptom onset, the four animals with earliest symptoms (28.5–34.9 h) had the lowest LF, LTx and EF levels at symptom onset. PA was <LOD in all four of these animals, ETx was <LOD in three of these animals, and bacteremia was negative in two and a low positive (+) in one. In these four animals with earliest symptoms, LF ranged from 0.076–1.3 ng/mL. The next sample, 6 hours later, showed that LF increased 5- to 13-fold to 0.538–6.4 ng/mL. However, PA and ETx remained <LOD, 6-hours later, in the two animals with LF levels below 1 ng/mL.

**Table 4.** Symptom onset biomarker levels.

| Animal ID | Hours to term | Symptom Onset-h | Sample time-h | Bact (next/term*) | Total-LF (next) | LTx | Total-EF | ETx (next) | Total-PA (next) | PA83 |
| --- | --- | --- | --- | --- | --- | --- | --- | --- | --- | --- |
| 1) C58282 | 48.5 | <u>29.1</u> | 30 | pos (5.3E+02) | 0.235 (2.99) | 0.103 | 0.00068 | <LOD (0.0045) | <LOD (9.11) | <LOD |
| 2) C59621 | 48.9 | 47.4 | 42~ | 2.9E+06 (1.3E+09*) | 57.8 (NA) | 23.7 | 20.7 | 1.87 | 333 (NA) | 67.4 |
| 3) C60763 | 54.7 | 54.7 | 48~ | 5.4E+06 (6.2E+08*) | 366 (NA) | 203 | 74.7 | 15.7 | 2222 (NA) | 259.4 |
| 4) C58176 | 55.2 | 38.2 | 42 | 9.1E+04 (3.1E+06) | 190 (NA) | 63.9 | 40.6 | 10.3 | 199 (NA) | 47.3 |
| 5) C58003 | 57.3 | 40.3 | 42 | 5.9E+06 (2.6E+05) | 220 (NA) | 97.8 | 6.89 | 2.0 | 1143 (NA) | <LOD |
| 6) C59656 | 57.9 | 40.9 | 42 | 7.0E+03 (3.6E+06) | 8.5 (75.2) | 0.893 | 0.059 | 0.00082 | 13.6 (731) | <LOD |
| 7) C58214 | 58.4 | <u>28.5</u> | 30 | neg (3.0E+02) | 0.136 (0.839) | 0.058 | 0.0005 | <LOD (<LOD) | <LOD (<LOD) | <LOD |
| 8) C60732 | 59.0 | 35.8 | 36 | 7.4E+06 (2.9E+03) | 313 (557) | 165 | 39.2 | 7.1 | 1189 (693) | <LOD |
| 9) C59669 | 64.8 | 56.3 | 60 | 1.9E+05 (4.0E+08*) | 200 (NA) | 149 | 5.62 | 0.872 | 282 (NA) | <LOD |
| 10) C57161 | 66.9 | 41.5 | 48 | 7.2E+06 (1.1E+06) | 366 (741) | 245 | 53.4 | 15 | 2221 (2303) | 516 |
| 11) C58170 | 74.1 | 61.4 | 60~ | 6.3E+04 (8.3E+08*) | 189 (NA) | 139 | 0.194 | 0.034 | 428 (NA) | <LOD |
| Mean-h<br>Median Fast | 58.7±7.6 | 43.1±10.8 | 41.3±9.9 | 1.9E+05 (3.5E+04–<br>5.7E+06) | 190 (33.2–<br>267) | 97.8 (12.3–<br>157) | 6.9 (0.13–<br>40) | 1.9 (0.017–8.7) | 333 (106–<br>1166) | ND |
| 12) C59619 | 76.4 | 35.3 | 36 | 1.8E+05 (1.5E+04) | 32.5 (37.4) | 12.5 | 0.661 | 0.278 | 368 (367) | 36.2 |
| 13) C58135 | 90.4 | <u>34.9</u> | 36 | 2.5E+02 (5.3E+03) | 1.30 (6.4) | 0.256 | 0.0053 | 0.00035 | <LOD (7.21) | <LOD |
| 14) C57793 | 91.1 | 67.0 | 72 | 9.2E+04 (1.2E+08) | 33.3 (7996) | 15.8 | 0.933 | 0.272 | 275 (34638) | <LOD |
| 15) C58167 | 91.7 | 37.3 | 42 | 8.5E+03 (7.1E+03) | 18.9 (19.4) | 6.49 | 0.38 | 0.082 | 129 (65.0) | <LOD |
| 16) C57998 | 100.8 | 77.2 | 84 | 2.9E+04 (3.1E+07) | 22.6 (1754) | 9.59 | 0.459 | 0.035 | 144 (15503) | 5.565 |
| 17) C59617 | 105.1 | 88.1 | 84 | 9.6E+03 (5.9E+04) | 93.8 (178) | 64.1 | 1.95 | 0.511 | 599 (813) | 25.4 |
| 18) C58159 | 109.0 | <u>30.2</u> | 30 | neg (pos) | 0.076 (0.538) | 0.029 | 0.00056 | <LOD (<LOD) | <LOD (<LOD) | <LOD |
| 19) C58122 | 115.1 | 35.6 | 36 | 8.4E+03 (3.6E+03) | 5.33 (22.0) | 1.29 | 0.044 | 0.00042 | 18.7 (152) | <LOD |
| 20) C57997 | 125.9 | 53.9 | 60 | 3.7E+02 (5.3E+02) | 15.5 (10.4) | 6.97 | 0.048 | 0.014 | 200 (96.7) | <LOD |
| 21) C58156 | 140.1 | 60.7 | 60 | 1.3E+04 (2.6E+02) | 19.8 (11.7) | 8.68 | 0.076 | 0.014 | 172 (137) | <LOD |
| 22) C59618 | 201.9 | 57.0 | 60 | 9.2E+02 (pos) | 8.51 (6.3) | 1.05 | 0.021 | 0.00065 | 25.7 (28.3) | <LOD |
| Mean-h<br>Median Slow | 113±34.4 | 52.5±19.5 | 54.5±19.8 | 8.5E+03 (6.5E+02–<br>2.1E+04) | 18.9 (6.9–<br>27.6) | 7.0 (1.2–<br>11.0) | 0.076<br>(0.033–0.56) | 0.014<br>(0.00054–0.18) | 144 (22.2–238) | ND |
| 23) C58174 | EOS | 108.4 | 108 | 3.4E+02 (neg) | 7.3 (4.2) | 1.6 | 0.041 | 0.0058 | 43.8 | <LOD |
| Mean-h 22-NS<br>Median All 23 | 86.1±37.1 | 47.8±16.1 | 48.9±17.4 | 1.3E+04 (6.5E+02–<br>1.9E+05) | 22.6 (7.9–<br>190) | 9.6 (1.2–<br>81.0) | 0.38 (0.043–<br>6.3) | 0.035<br>(0.00074–1.4) | 199 (22.2–398) |  |

#### 2.3 Kinetics of toxemia and bacteremia

##### Variable Kinetics of LF and bacteremia

During anthrax, vegetative *B. anthracis* and the toxins are found in circulation at varying levels at varying times. Bacteremia and LF represent the two sides of anthrax, infection, and intoxication. Bacteremia and LF graphed over time in all 23 animals demonstrate the variability of progression and the differences in these two biomarkers (Fig 2A,B). Most, but not all, of the animals appeared to have a triphasic kinetics of bacteremia and LF, with an initial rapid increase (phase-1), a plateau or decrease (phase-2), and a rapid increase leading to terminal stage (phase-3). The kinetics of bacteremia and LF were dramatically different in the 11 animals that progressed faster and died earlier compared to the 11 with longer survival times and slower progression. Specifically, early levels of bacteremia and LF appeared to be much higher in animals with faster progression and shorter survival than in the 11 animals with longer survival, and the one animal that survived. Overall, the kinetic trends, rises and declines, were similar between bacteremia and LF, but the phase-2 declines were more dramatic, declining for bacteremia compared to a plateau for LF.

**Figure 2.**
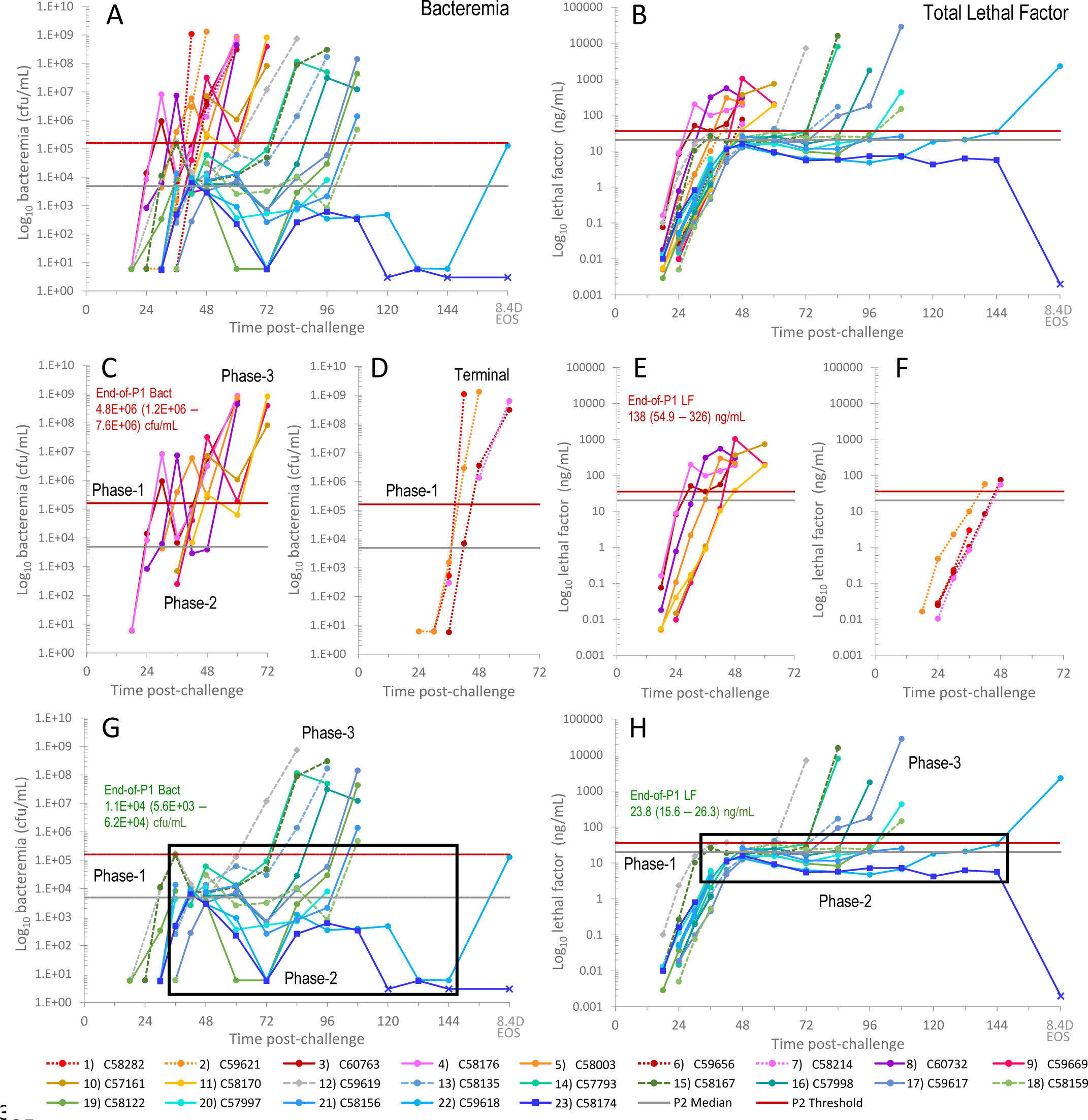
Kinetics of bacteremia and lethal factor in 23 cynomolgus macaques with variable progression. Bacteremia (Bact) in (cfu/mL) (A) and total lethal factor (LF) in (ng/mL) (B) over the course of infection in 23 cynomolgus macaques with inhalation anthrax. 11 animals had higher early bacteremia/toxemia and shorter survival (fast progression) (red/orange/purple lines), and 11 animals had lower early toxemia and longer survival (slow progression) (aqua/green lines), and one survivor with low early toxemia (blue line, square symbols). Most animals had triphasic kinetics consisting of an initial rise (phase-1), plateau or decline (phase-2), and final rise to terminal (phase-3). Bacteremia and LF in 7 animals with fast triphasic kinetics (C) and (E) and 4 animals with monophasic kinetics, a single rapid rise, (D) and (F), respectively. Bacteremia and LF in 11 animals with slow progression and survivor, (G) and (H), respectively. The last time point for 22) C59618 that died at 8.4 days (8.4D) and the survivor, 23) C58174, which declined to less than the limit of detection (LOD) at the end-of-study (EOS, 14- days/336 hours) are not to scale. For bacteremia (bact), non-quantifiable results of low positive were assigned a value of 6 and negative at 3 cfu/mL. Bacteremia and LF below LOD (x-symbol). Boxes represent the broad range of bacteremia in phase-2 (G) and narrow range of LF levels during phase-2 (H).

##### Disease progression

So, with higher early bacteremia and LF for the 11 animals with shorter survival time (faster progressing) and lower levels for the 11 animals with longer survival times (slower progressing), these early biomarker levels may distinguish subsequent progression. Separating these two groups, we show that of the 11 faster progressing animals, 7 had a triphasic kinetics of bacteremia and LF (Fig 2C, E), and 4 had monophasic progression, where bacteremia and LF appeared to have a single stage that increased directly from detection to terminal time (stage-3), unbroken by a plateau/decline (Fig 2D, F). The speed of progression meant that terminal plasma samples were not obtained for LF (or any toxin measurement) (Fig 2E, F). However, terminal measurements for bacteremia were available and provided evidence of the final rise to terminal (Fig 2 C, D). In animals with fast progression, end-of-phase-1 bacteremia and LF levels were significantly higher than in slow progression and the phase 2 plateaus were short in length, only 6 to 12 hours, compared to 24 hours and longer for animals with slow progression.

In the 11 non-survivors with slower progression, almost all had a triphasic kinetics of bacteremia and LF (Fig 2 G,H). In phase-1, LF increased to lower levels at the end-of-phase-1 (Fig 2H). In these animals, phase-2 provided a stable feature, with LF levels that varied by less than one order of magnitude from lowest at 4.8 ng/mL to highest at 41.5 ng/mL across all animals and time points (boxed area) (Fig 2H).

Conversely, phase-2 bacteremia varied over 4-orders of magnitude, from 1.76E+05 cfu/mL to low ‘+’ and was even non-detectable at several time points in the animal that survived to the end-of-study (Fig 2G). For other toxins, phase-2 for PA and LTx had less variability (similar to LF) and EF and ETx had more variability (similar to bacteremia) (S3 Fig). The final phase-3 rise in LF and bacteremia was rapid and to high levels for most animals. Those that did not reach high terminal bacteremia of 10^7^-10^9^ cfu/mL, or high LF, were either euthanized, for which timing may vary, or were determined to have died from meningitis rather than sepsis.

The progression of toxemia and bacteremia for the one animal that survived to the end of the study and one animal that survived to 8.4 days were similar for bacteremia out to 144 hours and LF out to 108 hours. Both animals had the lowest but positive bacteremia in phase-2, but the survivor had some negative points later (Fig 2G). The survivor resisted the pressures of toxemia and infection, whereas in the non-survivor, resistance was overcome.

##### All toxins and bacteremia: kinetics in individual animals

A complete list of all toxin levels for each animal at each time point is included in a separate file (S1 File) and bacteremia is included in S2 File. All toxins and bacteremia were graphed together for all 23 individual animals, numbered by survival time (Fig 3). Due to the speed of progression and hematological changes in the latest hours of anthrax, terminal plasma samples for toxin measurements were not available in 17 animals, all 11 with short survival and 6 with longer survival (Fig 3, animals 1-16, 19). However, whole blood for bacteremia was collected at terminal stage, allowing visualization of kinetic trends in the final hours of progression.

**Figure 3.**
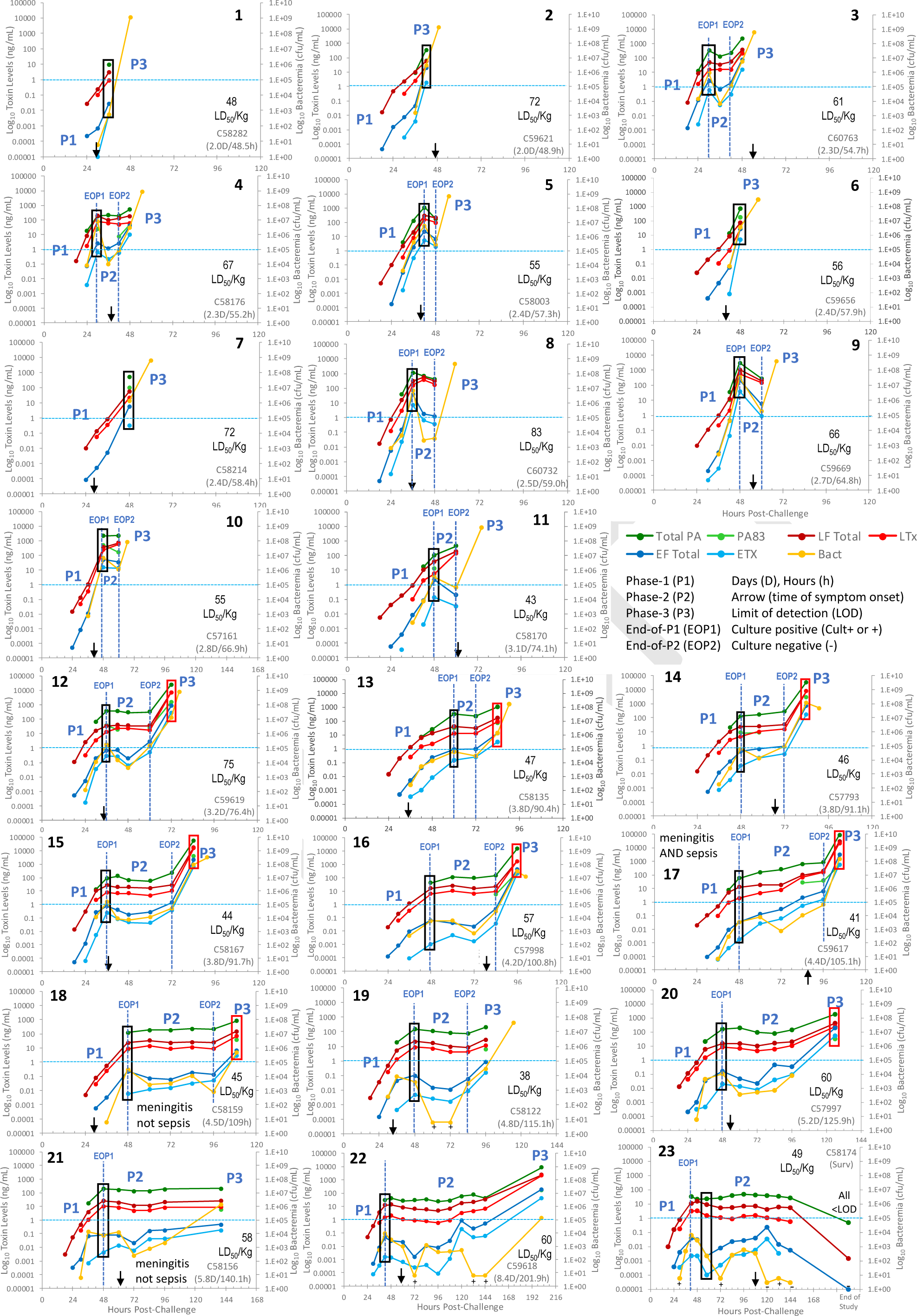
Anthrax toxins and bacteremia during inhalation anthrax in individual cynomolgus macaques. Total PA, PA83, LF, EF, LTx and ETx (ng/mL) and bacteremia (cfu/mL) in 11 animals with fast progression (numbered 1-11), 11 with slow progression (12–22), and the survivor (23). Phase-1 (P1), phase-2 (P2) and phase-3 (P3) are indicated when present. Blue vertical dashed lines indicate the point separating the phases; the end-of-P1 (EOP1) that ends the P1 rise and begins the P2 plateau and the end-of-P2 (EOP2) that ends P2 and begins P3. Black arrows indicate the time of symptom onset. Survival time-days (D), hours (h). Black rectangles at the end-of-phase-1 show how toxin levels are convergent (lower differences between highest toxin PA and lowest toxin ETx) in animals with shorter survival or divergent (higher differences between PA and ETx) in animals with longer survival. Red rectangles in phase-3 in animals with longer survival show convergent levels at terminal infection. Though units for toxins (ng/mL) and bacteremia (cfu/mL) are not directly comparable, both were graphed on a similar scale (10- orders of magnitude) for comparison of dynamic changes, orders of magnitude. Aqua horizontal dashed line at 1 ng/mL toxin concentration and 1.0E+05 cfu/mL bacteremia. Dynamic changes of bacteremia trended most closely with EF and ETx. Low positive culture (+) graphed at 6 cfu/mL and negative culture (-) at 3 cfu/mL. Less than the limit of detection (<LOD).

The toxin/bacteremia profiles of the 11 animals with shortest survival (fast progression) included four with monophasic progression with an initial rise in phase-1 and a continued rise in bacteremia at terminal stage (Fig 3, animals 1, 2, 6, 7). Animal 1 only had toxins measured out to 36 hours post-challenge and therefore, didn’t reach levels as high as the other three monophasic animals with toxins measured at 42 hours (animal 2) and 48 hours (animals 6, 7). Terminal stage bacteremia measured after the last toxin samples, reached high levels of 10^8^ to more than 10^9^ cfu/mL. In the other seven animals with fast progression, toxin levels only showed the phase-1 increase and a short phase-2 plateau/decline (6 to 12 hours). However, a third stage was evident with the phase-3 increase in bacteremia, also to high levels (animals 3-5, 8-11).

Kinetic profiles for the 11 non-survivors with longer survival (slow progression) were triphasic with one exception discussed below. Three phases were observed for all toxins, even in the six animals missing terminal toxemia (Fig 3, animals 12-16, 19). These 6 showed the phase-1 increase, the phase-2 plateau, and the final rise in phase-3 approaching terminal. The changes in bacteremia in the final few hours showed a continued rise in four (animals 12, 13, 15, 19), and a decline in two (animals 14, 16). For these two, changes in the internal environment, hypoxia, or other conditions near the terminal stage may have inhibited further replication of the bacilli. Animal 20 had a terminal low positive (+) bacteremia in the presence of high toxemia and was excluded. It was presumed to be a failed test since all other animals with high toxemia had high bacteremia.

Overall, trends were consistent in all animals. The detection of PA83 was sporadic indicating that PA63 was the predominant form of PA circulating in cynomolgus macaques. The kinetic trends were similar for bacteremia and all toxins, LF, LTx, EF, ETx, and PA. However, phase-2 declines in EF and ETx were more dramatic, similar to declines in bacteremia, whereas PA, LF, and LTx tended to plateau rather than decline in phase-2. The similar trends occurred whether the kinetics were monophasic with one continuous rise, or triphasic, with the rise-plateau/decline-rise. All toxins followed the same general pattern, with two exceptions. Animal 11 had a mixed triphasic kinetics with phase-2 declines for EF, ETx and bacteremia, while PA, LF, and LTx levels did not appear to plateau but continued to rise, albeit more slowly than in phase-1 (Fig 3, 11). Animal 21 had a long phase-2 plateau to terminal for all toxins, while only bacteremia started to rise in phase-3. The pathological cause of death in animal 21 was meningitis rather than sepsis.

##### Potential relationship with survival time

As seen above, LF and bacteremia rose to higher levels early in the 11 animals with fast progression. As ordered by survival time, there appeared to be an association between early toxin levels and subsequent survival time. Early toxin levels and bacteremia, for example, at the end-of-phase-1 (black box), were higher in animals with shorter survival and lower in those that survived longer. Using the horizontal line at 1 ng/mL toxin concentration and 1.0E+05 cfu/mL bacteremia as a gauge, you can see that at the end-of-phase-1 (first rise), toxins and bacteremia trended lower with longer survival. Another trend was the magnitude and length of the phase-2 plateau, with higher concentrations and shorter duration in animals with short survival and lower concentrations and longer duration with longer survival. This indicated that early toxin levels appeared to be associated with subsequent survival time. This association was explored in detail below.

##### Relationship of toxins to each other

In every animal, total PA, once positive, was the most abundant toxin, with PA>LF>LTx>EF>ETx (Fig 3). In animals with longer survival, the ranges of these toxins during phase-2 were more divergent, with 4- to 5-orders of magnitude between the concentrations of PA (highest) and ETx (lowest) and most divergent in animals 16-23 (Fig 3, black boxes). In phase-3, all toxins increased rapidly and converged (Fig 3, red boxes). In contrast, in animals with shorter survival, toxins reached higher levels and converged much earlier, approaching or at the end-of-phase-1 (Fig 3, animals 2-11). Lower toxin levels were associated with overall divergent toxin levels and high toxin ratios and higher toxemia with convergent toxin levels and lower toxin ratios.

### 2.4 Characterization of toxins, bacteremia, and toxin ratios stratified by phase as defined by toxemic profiles

#### 2.4.1 Phase-1

##### Toxemia in the early Incubation period

The earliest time points at 18- and 24-hours post-challenge were collected before earliest symptoms at 29 hours post-challenge. LF was positive in 12 animals by 18 hours post-challenge and in all 23 by 24 hours. Combined 18-to-24-hour median (interquartile range-IQR) LF was 0.025 (0.013–0.115) ng/mL (n=35). EF was positive in 5 animals at 18 hours and 16 at 24 hours, at 0.00021 (0.0014-0.00007) ng/mL (n=21). Other toxins and bacteremia were not detected at these early time points in sufficient numbers for assessment. These early phase-1 levels were similar for animals with fast and slow progression (Fig 4C, Phase-1).

**Figure 4.**
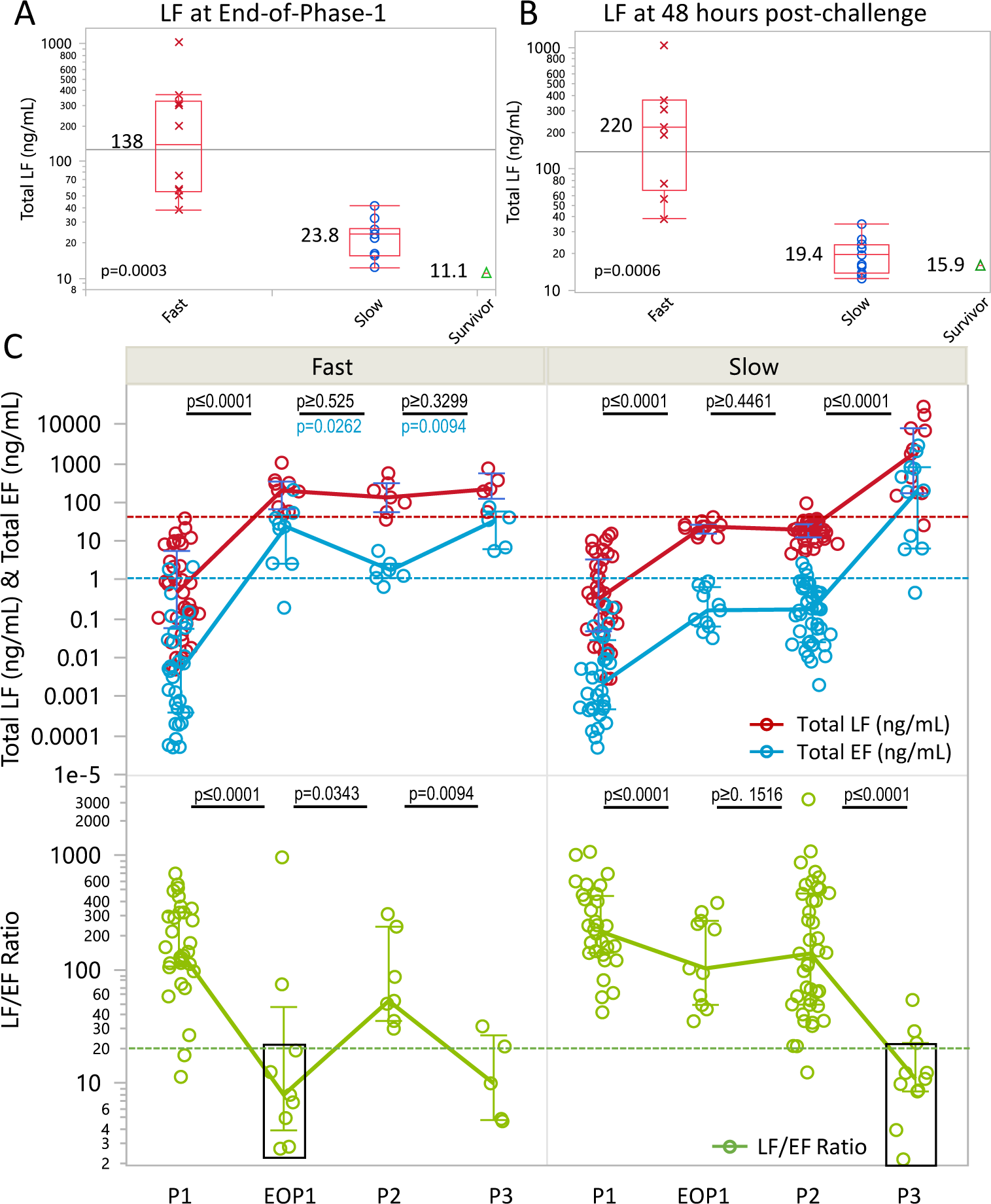
Stage-dependent biomarker levels. Lethal factor levels in animals with fast and slow progression, and survivor, at the end-of-phase-1 (A) and 48 hours (B). Box plot is for median and quartiles, error bars for minimum and maximums, excluding outliers, median levels and p-values included for Wilcoxon rank sum test. (C) LF, EF levels (upper panel), and LF/EF ratio (lower panel), during earliest detection in phase-1 (P1), at the end-of-phase-1 (EOP1), subsequent phase-2 (P2), and phase-3 (P3) for fast progression (left panel) and slow progression (right panel). Lines connect the medians and error bars give quartiles for each stage. Red dashed line represents the LF threshold and blue line the EF threshold for phase-2. P-values included for testing differences in toxin levels and toxin ratios between phases. Black font includes all parameters unless another color font is shown for a specific parameter. Black rectangle shows similarity of low LF/EF ratios at the EOP1 for fast progression and P3 for slow progression.

##### End-of-phase-1

The end-of-phase-1, also the beginning of the phase-2 plateau, was readily apparent in all but the four animals with monophasic progression (Fig 3). In these four, the phase-1 rise continues to terminal bacteremia, corresponding to the late-fulminant stage-3 (phase-3 without an observable phase-2). By the end-of-phase-1, all toxins and bacteremia, except for the unprocessed PA83, were positive in all animals. The timing to reach the end-of- phase-1 ranged from 30 to 60 hours post-challenge. Levels at the end-of-phase-1 were higher in the fast progression group, than in the slow progression group, and the survivor (Table 5 and Fig 4A). The differences were also observed at a specific time point near the end-of-phase-1, at 48h (Fig 4B). For either feature, the kinetically defined end-of-phase-1, or a timed collection at 48 h, higher toxemia at this early stage was associated with a faster course of infection and a shorter survival time. In addition to LF, end-of-phase-1 levels of all toxins and bacteremia were significantly higher in fast progression compared to slow (Table 5). Higher toxemia at the end-of-phase-1 in fast progression had lower toxin ratios and lower toxemia had higher ratios.

**Table 5.** Stage-dependent biomarker levels.

| Parameter | End-of-Phase-1 (Beginning-of-Phase-2) |  |  | All of Phase-2 |  |  |  | Phase-3 (≤12 hours from terminal) |  |
| --- | --- | --- | --- | --- | --- | --- | --- | --- | --- |
|  | Fast (n=10)<sup>10</sup> | Slow (n=11)<sup>11</sup> | Survived (n=1)<sup>1</sup> | Fast (n=7)<sup>15</sup> | Slow (n=11)<sup>52</sup> | Survived (n=1)<sup>10</sup> | Slow Phase-2 Threshold# (n=11)<sup>52</sup> | Fast (n=9)<sup>9</sup><br>7.2(6.8–10.2)-hours | Slow (n=10)<sup>11</sup><br>4.4(0–7.1)-hours |
| Total-PA (ng/mL) | 620 (328–1447) | 133 (54.2–200)<sup>c</sup> | 33.9 | 326 (199–1189) | 137 (67.4–205)<sup>a</sup> | 34.6 (26.2–41.3) | 342 | 509 (308–1477) | 8547 (813–34638)<sup>d</sup> |
| Total LF (ng/mL) | 138 (54.9–326) | 23.8 (15.6–26.3)<sup>b</sup> | 11.1 | 200 (98.1–366) | 20.2 (13.9–26.1)<sup>a</sup> | 6.7 (5.6–9.8) | 35.8 | 200 (66.5–337) | 1754 (171–7996)<sup>f (ns)</sup> |
| Lethal Toxin (ng/mL) | 54.7 (19.5–188) | 8.6 (4.6–10.3)<sup>b</sup> | 3.25 | 139 (50.9–245) | 7.38 (4.51–10.3)<sup>a</sup> | 1.3 (0.89–2.0) | 18.9 | 97.8 (25.7–192) | 1799 (92.9–7991)<sup>e (ns)</sup> |
| Total EF (ng/mL) | 22.2 (2.7–42.8) | 0.17 (0.064–0.66)<sup>a</sup> | 0.040 | 2.68 (1.8–34.9) | 0.172 (0.042–0.47)<sup>a</sup> | 0.025 (0.0074–0.040) | 1.10 | 20.7 (5.6–37.75) | 201 (6.5–797)<sup>e (ns)</sup> |
| Edema Toxin (ng/mL) | 3.4 (0.51–9.1) | 0.0057 (0.0020–0.15)<sup>b</sup> | 0.0023 | 0.663 (0.294–7.1) | 0.035 (0.0048–0.095)<sup>a</sup> | 0.0028 (0.0013–0.0046) | 0.312 | 2 (0.62–11.9) | 54.8 (2.9–272)<sup>d</sup> |
| Bact (cfu/mL) | 4.8E+06 (1.2E+06–7.6E+06) | 1.1E+04 (5.6E+03–6.2E+04)<sup>a</sup> | 6.4E+03 | 2.6E+05 (6.3E+04–7.2E+06) | 4.9E+03 (7.1E+02–1.4E+04)<sup>a</sup> | 2.4E+02 (5.3E+00–1.2E+03) | 1.6E+05 | 1.3E+06 (2.2E+05–3.4E+06)<br>0h-Term (n=11)<br>6.3E+08 (4.0E+08–8.8E+08) | 6.7E+06 (3.9E+05–9.7E+07)<sup>f (ns)</sup><br>0h-Term (n=10)<br>4.4E+07 (1.2E+06–2.1E+08)<sup>c</sup> |
| PA/LF Ratio | 4.8 (2.9–7.1) | 5.6 (3.4–7.9)<sup>g (ns)</sup> | 3.1 | 2.9 (1.4–3.8) | 7.0 (4.4–8.8)<sup>a</sup> | 5.3 (2.9–6.9) | 2.5 | 3.1 (1.4–7.6) | 4.3 (3.4–6.3)<sup>f (ns)</sup> |
| LF/LTx Ratio | 2.4 (1.8–2.9) | 2.7 (2.5–5.2)<sup>f (ns)</sup> | 3.4 | 1.8 (2.4–1.4) | 2.6 (2.3–3.8)<sup>b</sup> | 5.0 (4.4–6.6) | 1.7 | 2.2 (1.5–2.7) | 1.20 (1.0–1.8)<sup>d</sup> |
| LF/EF Ratio | 9.0 (4.4–18.2) | 103 (49.2–270)<sup>b</sup> | 278 | 31.9 (12.7–74.6) | 127 (50.8–400)<sup>b</sup> | 350 (234–761) | 21.4 | 10.1 (3.7–33.8) | 11.0 (8.6–22.8)<sup>g (ns)</sup> |
| EF/ETx Ratio | 5.4 (4.2–12.3) | 16.0 (6.0–30.7)<sup>e (ns)</sup> | 17.4 | 4.5 (3.6–5.7) | 5.5 (3.7–11.7)<sup>f (ns)</sup> | 9.9 (6.8–16.7) | 2.7 | 4.8 (3.5–8.8) | 3.7 (2.6–4.3)<sup>e (ns)</sup> |
| PA/ETx Ratio | 190 (148–615) | 19,733 (2144–31,667)<sup>c</sup> | 14739 | 323 (167–986) | 4565 (1383–23,958)<sup>a</sup> | 13,467 (8967–35804) | 506 | 171 (118–733) | 198 (86.6–338)<sup>g (ns)</sup> |

Differences between end-of-phase-1 ratios were significant for LF/EF and PA/ETx (Table 5). LF/LTx and EF/ETx trended lower for fast progression but the differences were not significant. PA/LF did not fit the trend and was similar for fast and slow progression at the end-of-phase-1. Individual animal measurements at the end-of phase-1 (or within 12 hours when later toxin samples were not available) for all toxins (including PA83), toxin ratios, and bacteremia, are included in S1 Table.

#### 2.4.2 Phase-2

The phase-2 plateau/decline varies in levels and length, ranging from 6 to 12 hours in faster progressing anthrax to more than four days in animals with slower progression (Fig 2 and 3). The end-of-phase-1 is also the beginning of phase-2. Therefore, the differences at the end-of-phase-1 continued during phase-2, though only briefly in fast-progression, with significantly higher levels of all toxins and bacteremia for the 7 animals with phase-2 in fast progression, compared to phase-2 in the 11 animals with slow progression (Table 5). Phase-2 toxin ratios were lower in fast progression compared to slow progression. The differences in all ratios were significant except for EF/ETx. In the one survivor (animal 23), phase-2 toxin levels and bacteremia were even lower and toxin ratios higher (except for PA/LF), than in animals with slow progression.

##### Phase-2 critical thresholds for entry to phase-3

In slow progression, phase-2 LF levels varied less than one order of magnitude for the entire phase-2 period in all 11 animals (Fig 2H). This represents a period of relative stability that seems to be sustained only over a narrow range of LF. As such, it also represents a period where relatively small increases in LF may move infection to the final stage 3.

Median, interquartile ranges and upper (95%) and lower (5%) confidence intervals characterize this stable period in slow progression. The upper 95% interval of phase-2 toxemia is like an upper confidence limit. Numbers above this limit fail the criteria for being classified or included in the phase-2 plateau (for these animals and this study). Therefore, the upper limit represents a potential threshold for phase-2, above which an animal has exited phase-2 and entered phase-3. For toxin ratios, the lower 95% confidence limit applies. The thresholds for all toxins and ratios are included (Table 5) and were used to determine the timing of entry to phase-3 below.

#### 2.4.3 Phase-3

Stage-3, late-fulminant anthrax, is the stage where cure is the most unlikely. The remaining survival time after reaching this stage can be as short as a few hours and usually no longer than one day (48).

Phase-3, as defined by toxin kinetics, is the final rise during the final hours of terminal infection as seen for each animal (Fig 3). Both toxemic phase-3 and clinical stage-3 represent the final hours of survival. Characterization of phase-3 requires consideration of two main points. One point is the speed of this stage which resulted in relatively few samples collected for toxin measurements and few at terminal times. Therefore, to characterize phase-3 for toxins which lack terminal samples, we chose to include any measurements for time points collected within 12 hours of terminal time. The second point is that for animals with fast progression, the kinetic features are compressed within the last 12-to-24 hours (Fig 3, animals 1-11). As such, for animals with fast progression, points within 12 hours of terminal may include those at the end-of-phase-1 or phase-2. Bacteremia, which did have terminal measurements, was characterized two ways; 1) with the same time points as available for toxins and 2) with only the final terminal time points (0 hours before terminal).

With the limitations described above, phase-3 toxemia was compared between animals with fast and slow progression (Table 5 and Figure 4C). Available phase-3 samples for fast progression were collected at 7.2 (6.8‒10.2) hours before terminal and earlier than those for slow progression at 4.4 (0‒ 7.1) hours before terminal. Phase-3 toxin levels and bacteremia were higher for animals with slow progression than fast progression for all points within 12 hours of terminal, but the differences were not significant except for PA and ETx (Table 5). Of the toxin ratios only LF/LTx was significantly lower in slow progression than fast progression. With rapid changes during this stage, these differences may simply be an artifact of the differences in collection times. To support this, we compared bacteremia at final terminal collection times, which was significantly higher in animals with fast progression than in those with slow progression (Table 5). Of the significant changes in ratios, LF/LTx had the lowest ratio in phase-3, approaching 1.0, indicating that LTx nearly equals total LF and 100% existing in complex with PA63.

Overall, for stage progression, all toxins and bacteremia increase from first detection to the terminal stage, whereas all toxin ratios (except PA/LF) declined from earlier stages to terminal phase-3. PA/LF was the only toxin ratio to deviate from this pattern. PA/LF did not change consistently through all the stages. These trends, increasing toxemia and declining ratios through the phases may be used to define early and intermediate stages, with the phase-2 thresholds indicating the transition to late-stage anthrax.

##### Comprehensive staging with anthrax progression

Figure 4C summarizes four aspects of these three phases, early phase-1, the end-of-phase-1, subsequent phase-2, and phase-3. LF, EF, and LF/EF ratios are representative of differences between phases and between fast and slow progression. In early phase-1, LF and EF levels, and LF/EF ratios, were similar between fast and slow progression with low levels and high ratios. From early phase-1 to the end-of-phase-1, there were significant increases in LF and EF levels and decreases in LF/EF ratios in both groups, but to higher levels and lower ratios in animals with fast progression. At the end-of-phase-1, most points in animals with fast progression were above the phase-2 thresholds indicated for LF and EF, whereas all points in animals with slow progression were at or below the thresholds. LF/EF ratios at the end-of-phase-1 in fast progression were low and similar to those at phase-3 in slow progression. For fast progression, LF levels did not change from the end-of-phase-1, through phase-2, and phase-3. Thresholds were already exceeded and only a short time followed. EF levels declined significantly from the end-of-phase-1 to phase-2, then increased from phase-2 to phase-3. The changes in EF between these phases also resulted in inverse changes in the LF/EF ratio. For slow progression LF and EF levels increased from early phase-1 to the end-of-phase-1, and LF/EF ratios declined significantly. There were no significant changes between the end-of-phase-1 and phase-2, whereas between phase-2 and phase-3 LF and EF levels increased, and LF/EF ratios declined significantly.

### 2.5 Relationship of toxemia and time

#### 2.5.1 Time to reach phase-2 threshold and time from threshold to terminal

The relationship between phase-2 toxin levels and the time to reach the toxin thresholds was calculated for each animal for all toxins and bacteremia. The survivor never even reached the phase-2 mean so was excluded. Animals with fast progression reached the thresholds at 38.3±7.4 hours and those with slow progression at 78.7±14.1 hours (Fig 5A). Thus, on average, the threshold was reached in half the time in the 11 animals that had shortest survival times compared to the 11 animals with longer survival times. Most of the animals in the fast progression group exceeded the phase-2 threshold by the end-of-phase-1 and levels of most toxins remained above it throughout their short phase 2 plateaus (when present). After reaching the threshold, the time to terminal was remarkably similar for all animals, with 21.0±7.3 hours for fast and 20.4±7.3 hours for slow progression. These calculations were consistent for all biomarkers. This suggests that the time to reach critical toxin thresholds is variable but once it is exceeded, the course is rapid and similar.

**Figure 5.**
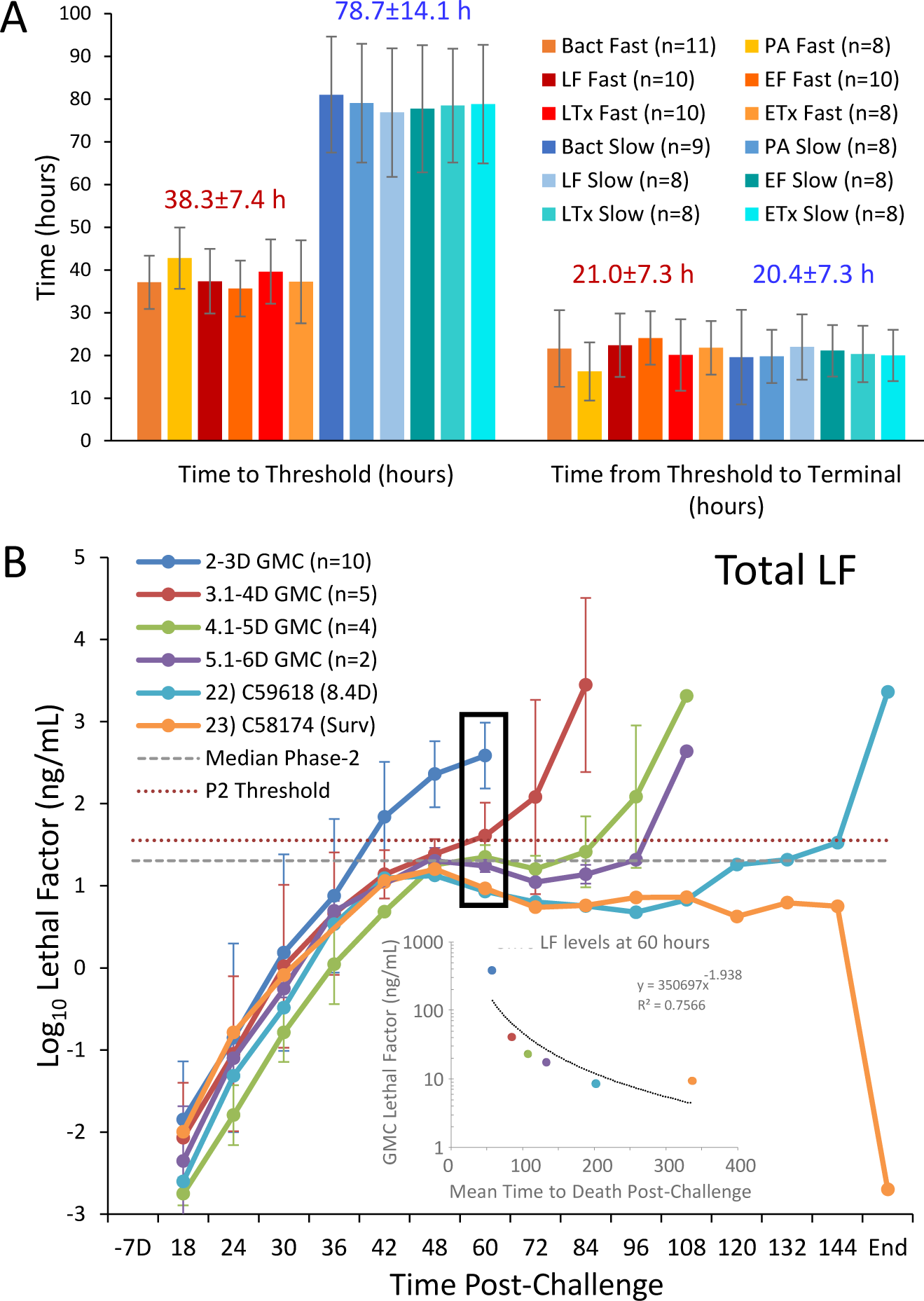
Relationship of toxemia and time. (A) Time in hours (h) from exposure to reach the phase-2 (P2)th threshold and time from threshold to terminal. The 95 percentile of phase-2 for all toxins and bacteremia in animals with slow progression represent an upper threshold for exiting phase-2 and entry to phase-3 (Table 5). The time to reach this threshold was calculated for individual animals and means for the number (n) animals for fast and slow progression. Error bars represent the standard deviation (SD). The overall mean±SD for all toxins and bacteremia combined for fast and slow progression is included above the bars. (B) Relationship between LF levels and survival time (days). Mean log_10_ concentrations and standard deviations as error bars for total LF over the time post-challenge (hours-h) for animals that died/euthanized at 1.9-3.0 days (n=10), 3.1 to 4.0 days (n=5), 4.1 to 5 days (n=4) and 5.1 to 6 days (n=2), and for one animal at 8.4 days and survivor to end-of-study (End). Final time points for the C59618 at 8.4 days and for C58174 the survivor at end-of-study (14 days) were included together (End). Horizontal lines shown for median phase-2 LF (dashed grey) and LF phase-2 threshold (red dotted line). The black rectangle for LF at 60 h showed mean log10 LF graphed for mean subsequent survival time (60-h to terminal) (inset), fitted with a power regression, indicating the inverse relationship of early LF levels with subsequent survival time.

#### 2.5.2 Relationship between LF levels and survival time

Our results for the fast and slow progression groups showed that animals with faster progression reached critical thresholds of toxin levels earlier and died sooner than those with slow progression. However, the time-to-death for animals with slow progression varied from 3.8- to 14-days. The relationship between LF levels and the time-dependent interval to death was demonstrated by grouping animals by survival time (Fig 5B). LF levels in animals surviving 1.9- to 3.0-days (n=10) reached the LF threshold first. Those surviving 3.1 to 4.0 days (n=5) reached the threshold second and the pattern continued with fewer animals and longer survival times reaching critical levels later. LF levels in the animal with the longest survival remained well below the phase-2 median until 108 hours, and in the animal that survived to the end of the study, LF levels remained below the phase-2 median throughout the entire course of infection. At 60 hours there was a negative relationship between LF levels and subsequent survival time. This relationship will be characterized in detail in the future. The pattern observed was strongest for LF but was present for all toxins and bacteremia (S4 Fig). Separation by survival time showed the relationship between toxin levels and survival time and the reliability with which exceeding the phase-2 threshold was followed by a short interval to death.

#### 2.5.3 Long progression of inhalation anthrax

The toxemia profiles for animal 23 that survived to the end of the study and animal 22 that survived 8.4 days were very similar with nearly identical for LF in these two animals out to 108 hours, both below the means for phase-2 (Figure 5B). Toxins were even lower in the non-survivor at 96 hours. By 108- to 120-hours toxins start to diverge. In the non-survivor. EF and ETx approached the phase-2 threshold at 120 hours, and LF by 144 hours (S4 Fig). PA and bacteremia remained below the mean in both animals out to 144 hours (the last scheduled sample before terminal). In the final sample, all toxins and bacteremia were non-detectable for the survivor and all toxin levels were high for the non-survivor.

### 3. Early hematology associated with time and progression

Samples were collected for hematological parameters through 72 hours in challenge group-1 and 144 hours in challenge group-2, prohibiting full comparison of hematology in all animals beyond 72 hours. Comparison of hematology was further limited to 48h because only two of 11 animals with fast progression had later measurements, prohibiting comparisons beyond 48h. All or most animals had measurements on day -7, 24-, 36- and 48-hours post-challenge, so these four time points were assessed for both changes with time and differences between fast (high early toxemia) and slow (low early toxemia) progression groups and the survivor. There were no combined effects of time and progression on early hematology. All hematology measurements are included in S3 File.

#### Changes in early hematology with time

Changes during early infection were significant (p ≤ 0.05) for many hematological parameters (Figure 6). P-values for time overall (TO) are included. There were significant increases through 48 hours in parameters associated with innate immunity, neutrophils, neutrophil%, neutrophil/lymphocyte (NL) ratio, NL% ratio, and white blood cells (WBC). Monocytes, monocytes%, large unstained cells, basophils, basophil%, eosinophils and eosinophil percentage increased at 24 hours and/or 36 hours, then declined at 36 hours and/or 48 hours. Lymphocytes and lymphocyte percentage declined from 36 hours to 48 hours. Parameters associated with red cells and oxygenation, hemoglobin, hematocrit, and red blood cells (RBC) declined significantly through 48 hours. These changes were observed for both fast and slow progression. Other parameters did not change significantly over time.

**Figure 6.**
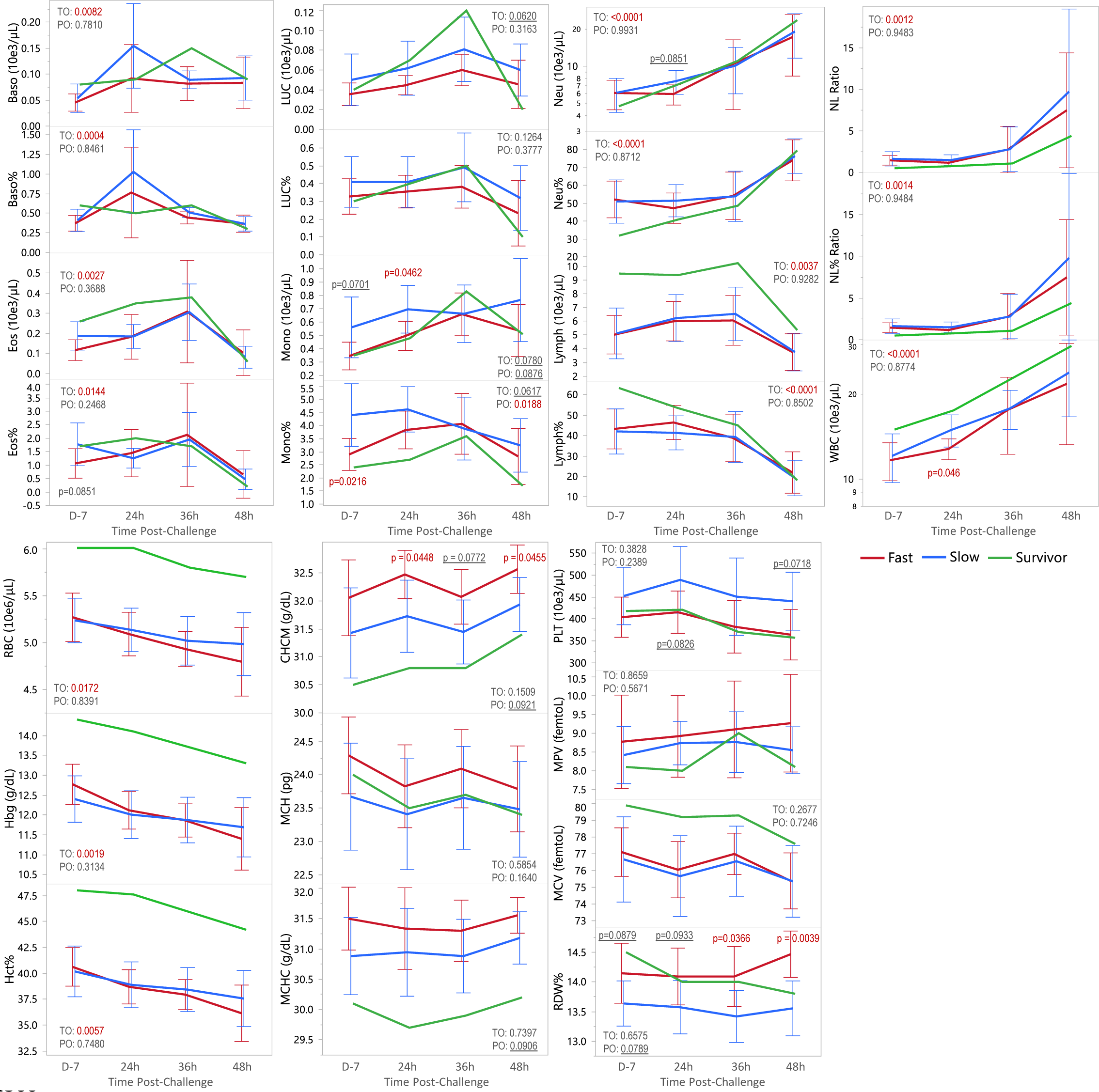
Early hematology during inhalation anthrax. Means and 95% confidence intervals graphed from Day -7 to 48 hours(h) post-challenge for fast progression (n=11, n=9 at 48 hours), slow progression (n=11), and survivor (n=1). Differences between animals with fast and slow progression at individual time points were determined by one-way ANOVA. P-values ≤0.10 were included in black font and p-values ≤0.05 in red font. All means, confidence intervals and p-values are included in Table 6. JMP fit model was used to test the effect of hematology with time (day -7, 24 hours-h, 36h and 48h) and progression (fast vs slow). P-value for time overall (TO) and progression overall (PO) is given. Basophils (Baso), eosinophils (Eos) large unstained cells (LUC), monocytes (Mono), neutrophils (Neu), lymphocytes (Lymph), neutrophil lymphocyte ratio (NL Ratio), white blood cells (WBC), red blood cells (RBC), hemoglobin (Hgb), hematocrit (Hct), corpuscular hemoglobin concentration mean (CHCM), mean corpuscular hemoglobin (MCH), mean corpuscular hemoglobin concentration (MCHC), platelets (PLT), mean platelet volume (MPV), mean corpuscular volume (MCV), red cell distribution width (RDW).

**Table 6.** Early hematology in cynomolgus macaques with inhalation anthrax.

| Parameter | Day -7 |  |  |  | 24 hours |  |  |  | 36 hours |  |  |  | 48 hours |  |  |  |
| --- | --- | --- | --- | --- | --- | --- | --- | --- | --- | --- | --- | --- | --- | --- | --- | --- |
|  | Fast (n=11) | Slow (n=11) | Surv | p-value | Fast (n=11) | Slow (n=11) | Surv | p-value | Fast (n=11) | Slow (n=11) | Surv | p-value | Fast (n=9) | Slow (n=11) | Surv | p-value |
| Baso<br>(10e3/μL) | 0.046 (0.024–0.067) | 0.054 (0.032–0.075) | 0.080 | 0.576 | 0.092 (0.023–0.161) | 0.155 (0.085–0.224) | 0.090 | 0.1965 | 0.082 (0.057–0.106) | 0.089 (0.065–0.114) | 0.15 | 0.6657 | 0.083 (0.039–0.128) | 0.093 (0.052–0.133) | 0.09 | 0.7476 |
| Baso% | 0.373 (0.258–0.488) | 0.409 (0.294–0.524) | 0.60 | 0.647 | 0.764 (0.243–1.28) | 1.03 (0.507–1.55) | 0.50 | 0.4639 | 0.446 (0.377–0.514) | 0.509 (0.441–0.577) | 0.60 | 0.185 | 0.367 (0.270–0.464) | 0.364 (0.276–0.451) | 0.30 | 0.9617 |
| Eos<br>(10e3/μL) | 0.117 (0.0595–0.175) | 0.188 (0.130–0.246) | 0.260 | <u>0.0851</u> | 0.184 (0.100–0.267) | 0.186 (0.102–0.269) | 0.35 | 0.9747 | 0.309 (0.116–0.502) | 0.306 (0.114–0.499) | 0.38 | 0.9835 | 0.104 (0.023–0.186) | 0.082 (0.008–0.155) | 0.06 | 0.6689 |
| Eos% | 1.06 (0.424–1.70) | 1.77 (1.13–2.41) | 1.70 | 0.118 | 1.45 (0.817–2.07) | 1.25 (0.626–1.88) | 2.00 | 0.659 | 2.13 (0.693–3.56) | 1.95 (0.521–3.39) | 1.7 | 0.8608 | 0.656 (0.042–1.27) | 0.473 (–0.082–1.03) | 0.20 | 0.6479 |
| LUC<br>(10e3/μL) | 0.036 (0.017–0.054) | 0.050 (0.031–0.069) | 0.040 | 0.270 | 0.045 (0.025–0.064) | 0.062 (0.043–0.081) | 0.070 | 0.1995 | 0.060 (0.036–0.084) | 0.081 (0.057–0.105) | 0.12 | 0.2132 | 0.046 (0.020–0.071) | 0.060 (0.037–0.083) | 0.02 | 0.3847 |
| LUC% | 0.327 (0.212–0.443) | 0.409 (0.294–0.524) | 0.30 | 0.307 | 0.355 (0.242–0.467) | 0.409 (0.297–0.521) | 0.40 | 0.4816 | 0.382 (0.231–0.532) | 0.491 (0.340–0.641) | 0.5 | 0.2978 | 0.233 (0.053–0.414) | 0.318 (0.155–0.481) | 0.10 | 0.4735 |
| Lymph<br>(10e3/μL) | 5.01 (3.48–6.54) | 5.11 (3.57–6.64) | 9.46 | 0.930 | 5.99 (4.52–7.47) | 6.22 (4.74–7.69) | 9.35 | 0.8239 | 6.05 (4.29–7.81) | 6.53 (4.77–8.29) | 10.2 | 0.6928 | 3.75 (2.41–5.09) | 3.75 (2.54–4.97) | 5.35 | 0.9948 |
| Lymph% | 43.2 (33.5–53.0) | 42.0 (32.2–51.8) | 63.3 | 0.857 | 46.4 (38.6–54.2) | 41.3 (33.5–49.2) | 53.4 | 0.3501 | 38.8 (27.6–50.1) | 39.3 (28.0–50.6) | 45 | 0.9532 | 21.9 (12.8–31.0) | 19.2 (10.9–27.5) | 18.1 | 0.6521 |
| Mono<br>(10e3/μL) | 0.346 (0.181–0.512) | 0.561 (0.396–0.726) | 0.350 | <u>0.0701</u> | 0.497 (0.359–0.635) | 0.696 (0.558–0.835) | 0.480 | **0.0462** | 0.660 (0.483–0.837) | 0.663 (0.486–0.840) | 0.83 | 0.9821 | 0.538 (0.268–0.807) | 0.766 (0.523–1.01) | 0.51 | 0.2028 |
| Mono% | 2.90 (2.00–3.79) | 4.41 (3.52–5.30) | 2.40 | **0.0216** | 3.84 (3.09–4.59) | 4.63 (3.88–5.38) | 2.70 | 0.1352 | 4.07 (2.97–5.18) | 3.89 (2.78–5.00) | 3.6 | 0.8112 | 2.82 (1.79–3.85) | 3.25 (2.31–4.18) | 1.7 | 0.5299 |
| Neu<br>(10e3/μL) | 6.09 (4.45–7.73) | 6.11 (4.47–7.76) | 4.76 | 0.982 | 5.97 (4.65–7.29) | 7.59 (6.27–8.91) | 7.15 | <u>0.0851</u> | 10.4 (5.62–15.2) | 10.1 (5.33–14.9) | 11.0 | 0.9281 | 17.2 (9.29–25.1) | 19.1 (11.9–26.2) | 23.53 | 0.7187 |
| Neu% | 52.1 (41.6–62.6) | 51.0 (40.4–61.5) | 31.8 | 0.874 | 47.2 (39.0–55.5) | 51.4 (43.1–59.6) | 40.9 | 0.4651 | 54.1 (41.3–67.0) | 53.9 (41.0–66.7) | 48.6 | 0.9762 | 74.0 (63.8–84.3) | 76.4 (67.1–85.7) | 79.6 | 0.7216 |
| NL Ratio | 1.46 (0.769–2.14) | 1.63 (0.947–2.32) | 0.503 | 0.707 | 1.17 (0.066–1.67) | 1.49 (0.986–1.99) | 0.765 | 0.3542 | 2.76 (0.19–5.33) | 2.80 (0.231–5.37) | 1.08 | 0.9813 | 7.48 (–1.32–16.3) | 9.74 (1.78–17.7) | 4.398 | 0.6937 |
| NL% Ratio | 1.45 (0.767–2.14) | 1.63 (0.947–2.32) | 0.502 | 0.704 | 1.17 (0.663–1.67) | 1.49 (0.986–1.99) | 0.766 | 0.355 | 2.76 (0.198–5.31) | 2.79 (0.236–5.35) | 1.08 | 0.9828 | 7.48 (–1.44–16.4) | 9.81 (1.75–17.9) | 4.398 | 0.6883 |
| PLT<br>(10e3/μL) | 404 (351–457) | 452 (399–505) | 418 | 0.192 | 415 (356–475) | 489 (430–549) | 421 | <u>0.0826</u> | 382 (311–453) | 451 (380–522) | 370 | 0.1687 | 364 (302–426) | 440 (384–497) | 357 | <u>0.0718</u> |
| MPV<br>(femtoL) | 8.77 (7.81–9.74) | 8.42 (7.46–9.38) | 8.10 | 0.593 | 8.92 (8.10–9.74) | 8.74 (7.92–9.55) | 8.00 | 0.746 | 9.1 (8.09–10.1) | 8.76 (7.76–9.77) | 9.0 | 0.6273 | 9.27 (8.34–10.2) | 8.55 (7.71–9.38) | 8.1 | 0.2407 |
| WBC<br>(10e3/μL) | 11.6 (9.69–13.6) | 12.1 (10.1–14.0) | 15.0 | 0.753 | 12.8 (11.3–14.3) | 14.9 (13.4–16.4) | 17.5 | **0.046** | 17.6 (13.6–21.6) | 17.8 (13.8–21.8) | 22.7 | 0.9387 | 21.7 (14.1–29.3) | 23.8 (16.9–30.7) | 29.56 | 0.6736 |
| CHCM (g/dL) | 32.1 (31.4–32.8) | 31.4 (30.7–32.1) | 30.5 | 0.199 | 32.5 (32.0–33.0) | 31.7 (31.2–32.2) | 30.8 | **0.0448** | 32.1 (31.6–32.6) | 31.4 (30.9–31.9) | 30.8 | <u>0.0772</u> | 32.6 (32.1–33.0) | 31.9 (31.5–43.2) | 31.4 | **0.0455** |
| Hgb (g/dL) | 12.8 (12.3–13.3) | 12.4 (11.9–12.9) | 14.4 | 0.294 | 12.1 (11.6–12.6) | 12.0 (11.5–12.5) | 14.1 | 0.753 | 11.9 (11.4–12.3) | 11.9 (11.4–12.3) | 13.7 | 0.9775 | 11.4 (10.7–12.2) | 11.7 (11.0–12.4) | 13.3 | 0.5534 |
| Hct% | 40.6 (38.6–42.7) | 40.2 (38.1–42.2) | 48.0 | 0.755 | 38.7 (36.9–40.5) | 38.9 (37.1–40.7) | 47.6 | 0.8791 | 37.9 (36.2–39.6) | 38.4 (36.7–40.1) | 45.9 | 0.6802 | 36.1 (33.5–38.8) | 37.6 (35.1–40.0) | 44.2 | 0.4188 |
| MCH (pg) | 24.3 (23.6–24.9) | 23.7 (23.0–24.3) | 24.0 | 0.178 | 23.8 (23.1–24.5) | 23.4 (22.7–24.1) | 23.5 | 0.379 | 24.1 (23.5–24.7) | 23.7 (23.0–24.3) | 23.7 | 0.3269 | 23.8 (23.1–24.5) | 23.5 (22.9–24.1) | 23.4 | 0.4909 |
| MCHC (g/dL) | 31.5 (31.0–32.0) | 30.9 (30.3–31.4) | 30.1 | 0.109 | 31.3 (30.7–32.0) | 30.9 (30.3–31.6) | 29.7 | 0.3892 | 31.3 (30.8–31.8) | 30.9 (30.4–31.4) | 29.9 | 0.2525 | 31.6 (31.2–31.9) | 31.2 (30.8–31.5) | 30.2 | 0.1435 |
| MCV<br>(femtoL) | 77.1 (75.2–79.1) | 76.7 (74.7–78.6) | 79.9 | 0.744 | 76.1 (74.1–78.0) | 75.7 (73.7–77.6) | 79.2 | 0.7752 | 77 (75.4–78.6) | 76.6 (75.0–78.2) | 79.3 | 0.6938 | 75.4 (73.4–77.3) | 75.4 (73.6–77.1) | 77.6 | 0.9842 |
| RBC<br>(10e6/μL) | 5.27 (5.04–5.50) | 5.24 (5.01–5.47) | 6.01 | 0.833 | 5.09 (4.88–5.31) | 5.14 (4.92–5.35) | 6.01 | 0.7655 | 4.93 (4.72–5.14) | 5.02 (4.81–5.23) | 5.8 | 0.5467 | 4.8 (4.45–5.14) | 4.98 (4.67–5.30) | 5.7 | 0.4047 |
| RDW% | 14.1 (13.7–14.6) | 13.6 (13.2–14.1) | 14.5 | <u>0.0879</u> | 14.1 (13.7–14.5) | 13.6 (13.1–14.0) | 14.0 | <u>0.0933</u> | 14.1 (13.6–14.5) | 13.4 (13.0–13.9) | 14.0 | **0.0366** | 14.5 (14.0–14.9) | 13.6 (13.2–13.9) | 13.8 | **0.0039** |

Means and 95% confidence intervals from Oneway ANOVA, assuming equal variances, for 11 animals with fast progression, 11 animals with slow progression, and one animal that survived to the end-of-study, on on day -7, 24-, 36-, and 48-hours post-challenge. P-values $\leq 0.10$ are underlined and p-values $\leq 0.05$ are in bold font. Basophils (Baso), eosinophils (Eos) large unstained cells (LUC), monocytes (Mono), neutrophils (Neu), lymphocytes (Lymph), neutrophil lymphocyte ratio (NL Ratio), white blood cells (WBC), red blood cells (RBC), hemoglobin (Hgb), hematocrit (Hct), corpuscular hemoglobin concentration mean (CHCM), mean corpuscular hemoglobin (MCH), mean corpuscular hemoglobin concentration (MCHC), platelets (PLT), mean platelet volume (MPV), mean corpuscular volume (MCV), red cell distribution width (RDW).

#### Differences between early hematology with progression

Considering the speed with which some animals hit the late stage, differences at any one or more early time points may influence progression. Differences in hematology parameters between animals with fast progression and slow progression were determined at individual time points (Table 6) and overall. P-values for significant parameters at individual time points, and all p-values for progression overall (PO) for the entire period are included (Fig 6). For progression overall, monocyte% was significantly higher in slow progression than fast progression (Fig 6). Conversely, red cell distribution width (RDW), corpuscular hemoglobin concentration mean (CHCM), and mean corpuscular hemoglobin concentration (MCHC) were higher in fast progression than slow, approaching significance (p ≤ 0.1) for the entire early period (p ≤ 0.0921) (Fig 6). These parameters and others were significantly higher at individual time points (Table 6 and Fig 6). Innate immune parameters were higher in animals with fast progression than slow; specifically, monocyte% on day -7, and monocyte count and white blood cells at 24 hours. Platelets were also higher in slow progression, approaching significance at 24 hours and 48 hours. CHCM was significantly higher in fast progression than slow progression at 24- and 48 hours and approached significance at 36 hours. RDW was significantly higher in fast progression than slow progression and 36 and 48 hours and approached significance at day -7 and 24 hours. Compared to all non-surviving animals, with both fast and slow progression, the survivor had higher lymphocytes, lower NL ratios, among the highest WBC count, exceptionally high RBC, hemoglobin, and hematocrit, higher MCV, and low MCHC and CHCM (Fig 6 and Table 6).

These results suggest that hematological differences may make for a more or less robust innate immune response (defense) or render the animal more or less susceptible to the destruction of red cells and associated physiologic consequences of anthrax (resilience). Collectively, the results suggest that higher levels of innate immune parameters (such as monocytes and WBC) at early time points (such as day -7 and 24 hours), were associated with slower progression and lower levels with fast progression.

Conversely, higher levels of parameters associated with disparately sized red blood cells, or hemolytic anemia (CHCM, MCHC and RDW) favored fast progression and lower ones slow.

### 4. Pathology

Assessment of terminal stage pathology can provide evidence of the impact of the toxemia and bacteremia measured during infection. Animals underwent complete necropsy, and gross and microscopic observations were recorded. Microscopic observations were assigned a semi-quantitative severity score as follows: minimal for a minimally detectable lesion (Grade 1), mild for an easily visualized lesion (Grade 2), moderate for a lesion affecting larger areas of the tissue (Grade 3) and marked for lesions affecting maximum areas of the tissue (Grade 4). Microscopic observations attributed to anthrax in this study included the presence of bacteria and other pathologic changes, such as fibrin, hemorrhage, necrosis, and inflammation (Table 7).

**Table 7.**
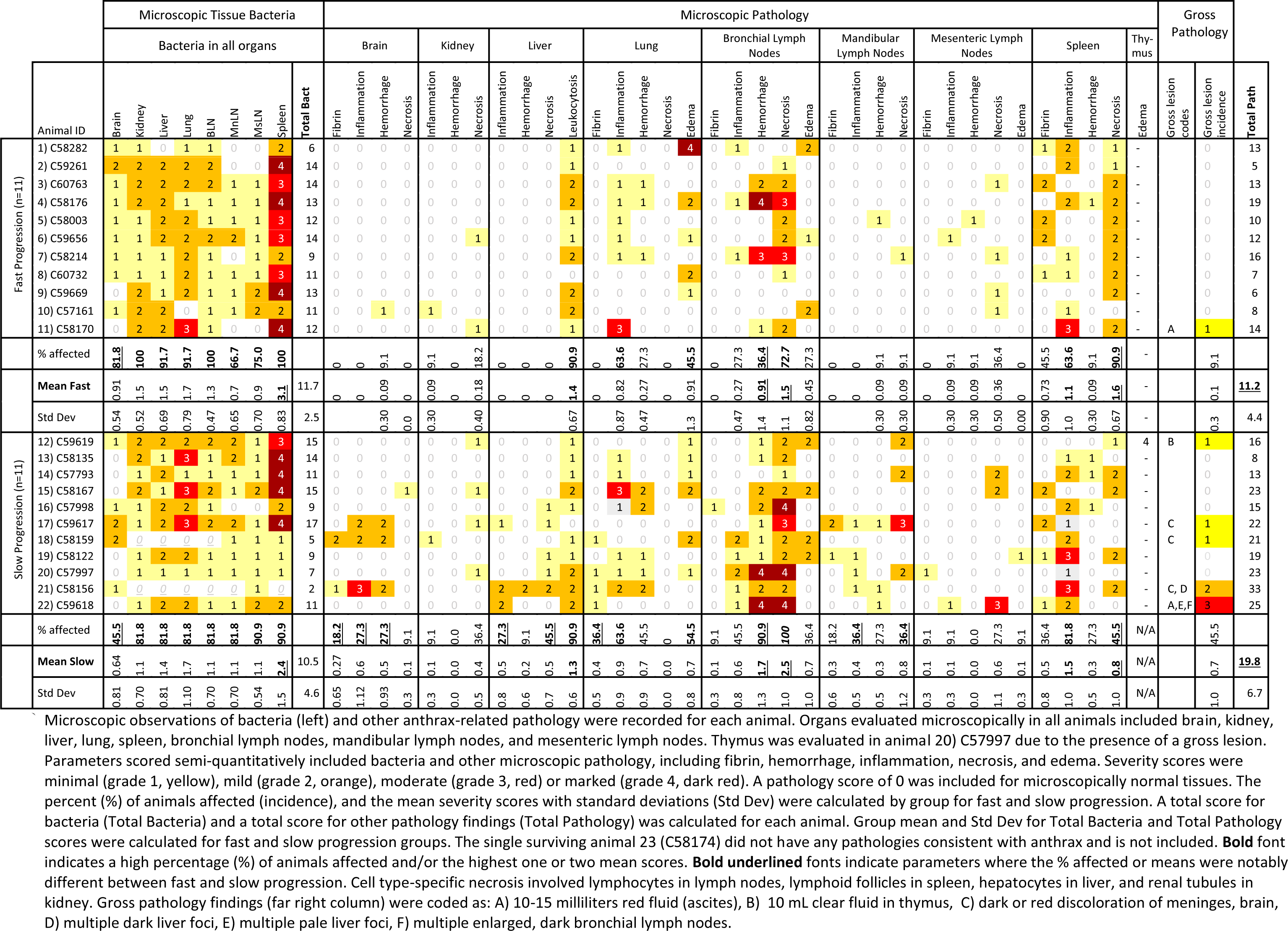
Microscopic and gross pathology summary.

#### Bacteremia and sepsis

Based on histopathological evaluation, both fast and slow progression animals had a high incidence (67 – 100%) of bacteria in organs and associated vascular spaces (Table 7). Overall, the spleen represented the greatest bacterial reservoir, with 21 animals affected and a severity grade of 3 or above in many of the animals (8 among fast progression and 5 among slow progression animals).

Thus, the highest group mean severity scores for bacteria were in the spleen, averaging 3.1 in the fast progression group and 2.4 in the slow progression group, followed by a mean severity score of 1.7 in the lungs in both groups. The only notable difference in bacterial tissue distribution between fast and slow progression animals was the presence of bacteria in brain in 81.8% of animals with fast progression and only 45.5% of animals with slow progression.

All but three of 23 animals in this study presented with features of fatal sepsis, as defined by *B. anthracis* dissemination to non-lymphoid organs including kidneys, liver, and lungs. Of the three animals without apparent sepsis, one (animal 23, C58174) survived *B. anthracis* challenge with no bacteria visible in any tissue. Two others, animals 18 (C58159) and 21 (C58156), were in the slow progression group and only had bacteria visible in the brain and lymphoid organs. Both animals had macroscopic observations of dark or red meninges at necropsy that correlated with microscopic fibrin, inflammation, and/or hemorrhage. Additionally, seizures were observed in animal 18 prior to euthanasia, and terminal bacteremia measurements in animal 18 and terminal toxin measurements in animal 21 were lower than those seen in animals with sepsis (Fig 3, animal 18, 21). Together, these findings supported that death in these two animals was a consequence of meningitis rather than sepsis. A third animal in the slow progression group, animal 17 (C59617), also had macroscopic and microscopic evidence of meningitis but with concurrent sepsis, disseminated bacteria to all organs, and high terminal toxemia (Fig 3, animal 17).

#### Histopathology

Among the tissues collected and processed for histopathological analysis, liver, lung, bronchial lymph nodes and spleen had highest incidence of anthrax-related microscopic lesions, including edema, fibrin, inflammation, hemorrhage, and necrosis (Table 7). The incidence and severity of microscopic changes in the kidney, liver, and lung were comparable between fast and slow progression animals. Lesions in other organs often had higher incidence and/or severity among slow progression than fast progression animals, including brain fibrin, inflammation, hemorrhage, and necrosis; liver inflammation, hemorrhage, and hepatocellular necrosis; bronchial lymph node edema, inflammation, hemorrhage, and lymphocyte necrosis; mandibular lymph node fibrin, inflammation, hemorrhage, and lymphocyte necrosis; mesenteric lymph node lymphocyte necrosis; and splenic inflammation. Among fast progression animals, a higher incidence and mean severity of splenic lymphoid necrosis was noted as compared to slow progression animals.

Despite most fast progression animals having bacteria visible in the brain, there was little associated tissue change (fibrin, inflammation, or necrosis) and only one animal with minimal hemorrhage. As previously noted, tissue changes in the brain of animals with slow progression were primarily limited to the three animals with meningitis, and meningitis was not seen in any animal that died less than 105 hours (4.4 days) post-challenge.

To facilitate global comparison between fast and slow progression animals, Total Bacteria scores were calculated for each animal by adding the individual microscopic severity scores for bacteria in all examined organs. Total Pathology scores were calculated for each animal by adding the individual microscopic severity scores for all anthrax-related lesions (except for bacteria) in all examined organs as well as the incidence of any anthrax-related gross lesions. Group mean (±standard deviation) Total Bacteria scores were 11.7±2.5 for fast progression and 10.5±4.6 for slow progression. Conversely, group mean Total Pathology scores were 11.2±4.4 for fast progression and 19.8±6.7 for slow progression, demonstrating higher pathology overall for animals with longer survival times and thus longer toxin exposure.

The single animal surviving to end-of-study (day 14) had non-specific pathology that included minimal chronic active inflammation in the lung and pale liver nodules, thought to be from a prior unrelated infection, with no microscopic evidence of anthrax.

## Discussion

### Anthrax progression

Historically, clinical inhalation anthrax was described as a biphasic disease, with two stages of ‘illness’, early-prodromal and late-fulminant (3, 48). The first stage lasted several days and was followed by either direct progression to the late-fulminant stage or a period of improvement before entry to the late stage. During a 1957 outbreak, Plotkin et al described five inhalation anthrax cases where after initial symptoms, one patient continued to work and two of the patients’ conditions were briefly ‘improved’ (48). However, after onset of the final stage in four patients, they only survived 7 to 25 hours. The 2001 inhalation anthrax cases allowed refinement of the clinical description to include three stages: 1) the early-prodromal, 2) intermediate-progressive, and 3) the late-fulminant stage (7, 33, 34). The kinetics of anthrax toxins determined previously in five rhesus macaques showed that PA, LF, LTx, EF, and the poly-glutamic acid capsule kinetics were triphasic, with a rise, a plateau (or decline), and a final rise: potentially comparable to the three human clinical stages of anthrax (22, 23, 45, 46). This study, with 23 cynomolgus macaques, allowed the first determination of all major anthrax toxins collectively, along with quantitative bacteremia, through all stages of anthrax progression. The larger numbers of animals (cynomolgus macaques) in this study yielded variations in progression compared to that seen in rhesus macaques.

Two different patterns of toxemia emerged in the cynomolgus macaques in this study. In 11 animals the toxin levels and bacteremia reached higher levels early, at 30-to-48 hours, compared to the other 12 animals in the study, which had much lower levels during this period. The 11 animals with high early toxemia/bacteremia happened to have the fastest disease progression and earliest deaths. In contrast, the animals with lower early toxemia survived longer. Thus, the kinetics defined two distinct survival groups, one with high early toxemia and fast progression and one with lower early toxemia and slow progression. This allowed characterization and comparison of both types of infection progression in a way that has not been done before. Some differences might be related to the higher spore dose per body weight in fast progression, but the differences were not statically significant. Furthermore, there was no significant correlation between LD_50_/kg and time-to-death, or LD50/kg and early toxin levels or bacteremia (data not shown). Nonetheless, there was a higher average LD_50_/kg for animals with fast progression. This prompts consideration, if feasible, for tailoring animal study design to target the spore dose according to an animals’ body weight.

Animals with slow progression exhibited a rise/plateau/rise, a triphasic kinetics, similar to the five rhesus macaques described previously. The triphasic kinetics of slow progression can be aligned with the 3 clinical stages (Fig 7), similar to that suggested previously (49). Phase-1 begins with early toxemia, LF and EF, appearing during the incubation period, before symptom onset. In the later part of phase-1, other toxins, LTx, ETx, PA, and bacteremia appear, along with early symptom onset in some animals.

**Figure 7.**
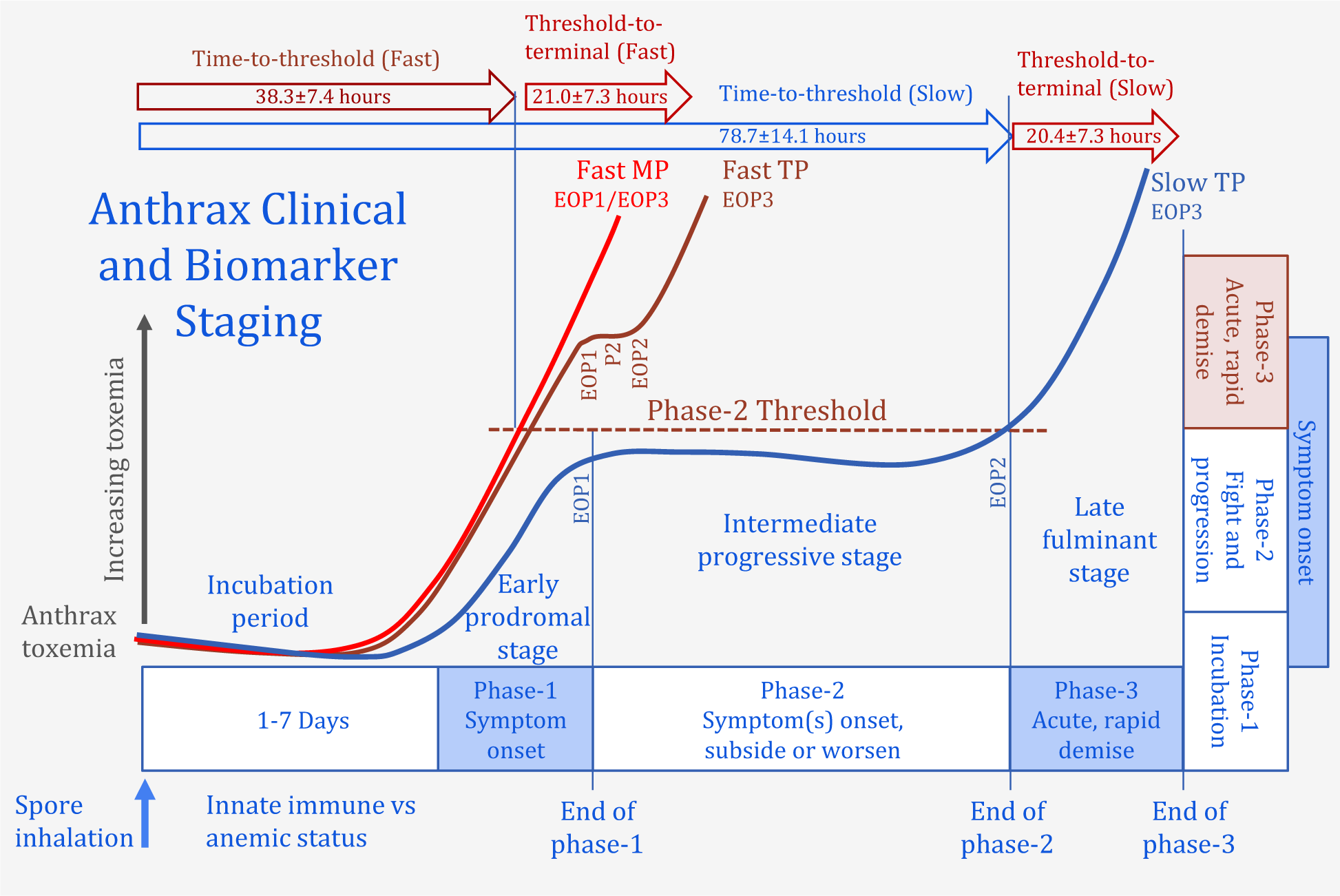
Proposed relationship of toxemia-defined progression with clinical staging. The triphasic (TP) kinetics of anthrax toxemia with slow progression in non-human primates mirrors the clinical staging algorithm. A brief incubation period is followed by 1) the early-prodromal stage 1 with the onset of non-specific symptoms which coincides with the early rise in toxemia to the end-of-phase-1 (EOP1). This is followed by 2) the intermediate- progressive stage which coincides with the plateau/decline in toxemia of variable duration that continues to the end-of-phase-2 (EOP2) when the critical toxemic thresholds of phase-2 are exceeded. Exceeding the threshold(s) coincides with entry to 3) the late-fulminant stage with a rapid decline in clinical stability and rapid rise in toxemia/bacteremia. In fast progression this occurs over a compressed timeline during which the phase-2 plateau may be observed (fast triphasic-TP) or may not be observed (fast monophasic-MP) kinetics. The dependence of progression on toxemia can be seen with the shorter time-to-threshold for fast progression than slow and the similar time from threshold-to-terminal for both. Progression may be reoriented from an apparent time/clinical- dependent staging to suggest possible toxin/clinical-dependent staging.

Toxins continue to rise to intermediate levels at the end-of-phase-1 (Table 5, median levels of 23.8 ng/mL for LF). This period is analogous to the early-prodromal clinical stage. The end-of-phase-1 begins phase-2 where toxins levels plateau (PA/LF/LTx) or decline (EF/ETx/bacteremia) for an extended period, analogous to the intermediate-progressive stage 2. LF levels throughout phase-2 were stable, ranging less than one order of magnitude over all time points and animals as described. Characterizing the entire stable phase-2 period allowed determination of the upper 95% confidence limits of this range, establishing a phase-2 threshold, above which phase-2 has ended and phase-3 commenced. After the plateau, toxins started to rise again and animals reached the phase-2 threshold (35.8 ng/mL for LF), marking entry to phase-3 with rapid accumulation of toxemia and rapid demise, corresponding with the late-fulminant stage.

For fast progression, the kinetics was predominantly triphasic with a rapid rise, shorter phase-2 plateau, and final rapid rise. A few animals appeared to be monophasic, with the phase-2 plateau skipped, or not discernable within the early 6-hour collection times of this study. The kinetics of all animals with fast progression occurred within a compressed time frame (Fig 7). As such, they reached higher levels at the end-of-phase-1 (Table 5, 138 ng/mL for LF), already above the phase-2 threshold of 35.8 ng/mL for LF and for all toxins and bacteremia, indicating early entry to phase-3. In monophasic progression the end-of-phase-1 and end-of-phase-3 are virtually synonymous. Passing the critical phase-2 toxin thresholds early led to rapid death, even before symptoms were observed in two of the animals.

An early study by Smith and Keppie identified a bacteremia threshold above which antibiotic treatment failed (11). This bacteremia threshold represents a point of no return beyond which an animal will not recover with antimicrobial treatment. In clinical anthrax the late-fulminant stage 3 has the lowest cure rate, 97% fatality for a century of inhalation anthrax cases (5). Thus, it is analogous to the point of no return. Stage-3 is analogous to phase-3 in this cynomolgus macaque model, indicating the phase-2 bacteremia/toxemia thresholds for entry to phase-3 may also predict stage-3 based treatment failures.

When toxemia went above the phase-2 threshold, toxins levels rapidly increased leading to the terminal phase. The mean time to reach the thresholds and enter phase-3 was on average, twice as long in slow progression as fast. In contrast, the mean time from threshold to terminal was nearly identical for both fast and slow progression at 20-21 hours, consistent with the duration of stage 3 described in clinical cases (3, 7, 48, 50). The findings support reorientation of the progression scheme from a time-dependent to a toxin-dependent one (Fig 7). Animals that rapidly pass the critical threshold in phase-1 show a compression (or elimination) of the intermediate-progressive phase-2. The 11 animals with the shortest survival times had an accelerated progression due to early high toxemia. Evaluation of toxemia by survival time supports the critical phase-2 threshold of toxin levels in the cynomolgus macaque model, since no matter the time post-challenge, once the threshold is passed, terminal stage-3 follows rapidly and within about 21 hours (Fig 5).

The fast progression observed in almost half of the animals is a challenging problem since it leaves little time for successful treatment after symptoms. A rapid course has been suggested by evidence from the Sverdlovsk outbreak for which dates of both symptom onset and death were available for 50 cases (51). The time from onset to death was on the same day for one case and within one day for 8 cases, consistent with the faster progression in this study. For all other cases, the time from symptom onset and death were 2 days and longer, which is more consistent with slow progression in our cynomolgus macaque study. A systematic review for a century of inhalation anthrax reported four cases with 1.5- to 1.6-days following initial symptoms (5). Though the incubation periods and exposure doses are not certain, the rapid death following onset suggest the possibility of rapid progression in humans following inhalation of *B. anthracis* spores, especially the weaponized spores in the Sverdlovsk cases.

### Other factors

Although, the average spore dose per kg was higher for fast progression than slow, there was no correlation of LD_50_/kg with early toxemia (data not shown). Vasconcelos *et al.* also observed ‘a tendency’ for cynomolgus macaques receiving higher spore doses to die more quickly than those that received lower doses (37). Early hematology showed trends related to time or progression. There were time-dependent changes over the first 48 hours, increases in innate immune cells, such as white blood cells and neutrophils, and declines in red blood cells, hemoglobin, and hematocrit. These occurred in both fast and slow progression, consistent with those seen in other animal models (52, 53). However, there were key differences between fast and slow progression for some parameters. Means for innate immune markers, especially, monocytes, were lower in fast progression than slow progression. Of course, this means that monocytes were higher in slow progression, perhaps offering more defense in these early stages. Red cell distribution width (RDW) and corpuscular hemoglobin concentration mean (CHCM) were higher in animals with fast progression and lower in animals with slow progression. High RDW is associated with deficiencies of iron, B-12, or folate and can be due to macrocytic anemia, underlying chronic inflammation, or infection (54). Studies in middle aged and older adults showed that even modestly elevated RDW was associated with an increased risk of all-cause mortality (55). Elevated RDW was also associated with increased severity of sepsis (56, 57).

The anthrax toxins actions make the differences in these factors even more important. LTx is known to cause anemia and has also been shown to suppress erythropoiesis, exacerbating the anemia, and promoting progression (58). Hemoglobin and hematocrit have been shown to drop during clinical inhalation anthrax and was observed here (59). In this study, the lone survivor, animal 23, had higher early white blood cells, notably lymphocytes, but also had exceptionally elevated red blood cell count, hemoglobin, and hematocrit (Fig 6). In contrast to animals with fast progression that had higher CHCM, the survivor also had the lowest CHCM (and MCHC) of all animals. These hematological markers will be further evaluated for their association with progression and outcome in future studies.

Two animals with long progression, animal 23 that survived and animal 22 that succumbed at 8.4 days, had very similar toxemic profiles, especially for LF. All toxins and bacteremia were well below the phase-2 mean out to 108 to 120 hours, but eventually diverged with toxins increasing in the non-survivor and declining to non-detectable in the survivor (Fig 5B and S4 Fig). There were some differences in these animals that might have contributed to the final outcome. The LD_50_ spore dose was higher in the survivor but when adjusted for weight was higher in the non-survivor (Table 1). Also, at 132 hours, the RBC count was within normal limits (5.0 to 7.6 e^6^/µL, Battelle report) at 5.24 e^6^/µL in the survivor but was below normal in the non-survivor (3.85 e^6^/µL). The higher beneficial hematological features described above for the survivor may have provided a better level of resilience against toxins and defense against the organism for recovery.

### Pathological factors associated with disease progression

Assessment of the degree of microscopic bacteria and other pathology at terminal-stage provides an assessment of the burden and impact of anthrax resulting from the varying toxemic progressions. The general findings were consistent with those described for cynomolgus macaques previously, with primary involvement in the spleen, bronchial lymph nodes, and lungs (37). Overall, animals with longer survival times and thus longer toxin exposure had a higher incidence and/or severity of pathologic findings but slightly lower severity of intralesional bacteria.

While both fast and slow progression groups had similar incidences of bacteria in the spleen, the greatest bacterial reservoir organ in this study, the mean severity of splenic bacteria was higher in animals with fast progression than slow progression. Splenic lymphoid necrosis was also seen in twice as many fast progression animals, and with twice the average severity, as compared to those with slow progression. Thus, both bacterial colonization and necrosis in the spleen were early to develop in the course of rapidly progressing anthrax. Conversely, microscopic lesions in other organs were often similar among all animals or had higher incidence and/or severity among slow progression animals. The latter category included bronchial lymph node lymphocyte necrosis and hemorrhage, suggesting that longer toxin exposure may allow for lesion development.

Although there were more fast progression animals with visible bacteria in the meninges than slow progression animals, they had little to no associated brain pathology. Studies have indicated that ETx plays a role in disrupting the blood brain barrier (BBB) function and LTx in promoting microbial BBB penetration (60). The higher early toxemia in fast progression may allow BBB penetration and penetration of bacilli, but death is rapid with little associated pathology. In animals with slow progression, the lower end-of-phase-1 and phase-2 toxemia may not facilitate penetration of the BBB quite as readily, but when bacteria did gain access to the brain, the longer low-level toxin exposure in phase-2 may have contributed to gross and microscopic fibrin accumulation, inflammation, and hemorrhage. These differences suggest the rapid sepsis of fast progression doesn’t allow sufficient time for a tissue response in the brain, whereas the longer course in slow progression allows for longer toxin exposure resulting in meningitis and death. Notably, meningitis was only seen in animals surviving at least 105 hours post-challenge. More may be learned from later studies in cynomolgus macaques, which have included the analysis of toxin levels in terminal organ tissues to help understand differences in the organ pathologies observed.

### Symptom onset biomarkers

LF, LTx, and EF were positive in all 23 animals at or before symptom onset, with LF the most abundant and EF the least abundant of these (Table 3). Although median PA levels at symptom onset were higher than LF, four animals were negative for PA. However, LTx (PA-LF) was positive (LOD of 0.0075 ng/mL) in these, indicating PA is present but below the PA detection limit (1.87 ng/mL) (Table 2). These four negative PA animals also had the earliest time of symptom onset of about 30 hours. Six hours later, LF increased 5- to 10-fold, leading to a conversion of PA positivity in two animals at low ng/mL levels, but not enough for all to be above the LOD for PA. The important conclusion is that we have defined the range of levels of multiple toxins at symptom onset in this high dose exposure model. These may help guide development of point of care diagnostics.

### Pre-symptom onset detection

Early diagnosis and treatment will improve the odds of surviving anthrax. With the analytical methods used in this study, LF was the best biomarker for pre-symptom onset detection. It was detected earliest and at higher levels than the next earliest detected biomarker, EF. In this study, LF and EF were detected before bacteremia and before symptom onset in every animal (Fig 1 and Table 3). We previously measured LF in human cutaneous anthrax in the absence of sepsis, and even after antibiotic treatment (61). In these cases, we presume that the LF detected is secreted locally from the site of infection, the cutaneous lesion, and is detected in circulation. Early detection of LF during inhalation anthrax, before bacteria are detectable in the blood, may originate from early events, such as germinating spores within macrophages (62).

### Toxemia in the absence of bacteremia

As shown at pre-symptom onset, anthrax toxins are often present in the serum in the absence of detectable bacteremia. In this study, LF and EF were detected before bacteremia in all of the cynomolgus macaques. In a previous study with 5 rhesus macaques, LF also tested positive before bacteremia, and it continued to test positive throughout the course of infection (46). In that rhesus macaque study, bacteremia went from positive - to negative (during phase 2) in some animals - back to positive, while LF was positive at all time points. As described above, LF was detected in serum of patients with cutaneous anthrax; after up to 7 days of antimicrobial treatment, and often when swabs of the cutaneous lesion were negative for tests detecting the bacterium (61).

Furthermore, LF was detected in serum and pleural fluid in two inhalation anthrax patients up to 11 days after initiation of antibiotic treatment when bacteria would no longer be detectable, and 5 days after antitoxin treatment. LF was also detected in an anthrax-like illness in a welder, caused by infection with *Bacillus tropicus* that produced anthrax toxins (in review). In this case, LF was detected in serum or plasma 11 days after antibiotics and in pleural fluid collected 38 days after antibiotics and two doses of antitoxin.

### Stage dependent toxemia and the phase-2 threshold

We identified toxemia associated with specific stages (Phase-1, -2, and -3). In the earliest stages of infection, detection was limited to LF and EF. The detection of other toxins and bacteremia came at later stages of phase-1 or early phase-2. The analytic sensitivity of the catalytic toxin methods allowed LTx and ETx to be detected before total-PA, though at lower levels than the total toxins, LF and EF. Both LTx and ETx comprise a fraction of the available PA63 and signal the point when active toxin complexes begin to accumulate. The appearance of LTx at 30.5 hours closely coincides with the earliest detection of bacteremia at 30.0 hours (including bacteremia that was below the LOQ and reported as a low ‘+’) (Table 3). The appearance of ETx at 33.4 hours coincides with the detection of low level but quantifiable bacteremia at 34.2 hours. This suggests that toxin complexes (LTx and ETx) are detectable in blood at the same time as bacteremia. PA was detected later, due in part to the higher detection limit, but also possibly due to the rate of cellular uptake of PA during early stages, resulting in less PA in circulation. Though first detection of total-PA was latest, once positive, it was the most abundant toxin; approximately 2.4-fold more abundant than LF at first positive PA. PA83 was only detected sporadically and was most consistently positive at high levels of total PA. This indicates that PA83 is rapidly hydrolyzed and activated to PA63, early and at all stages, in all cynomolgus macaques, as also seen in rhesus macaques (45). Therefore, PA83 detection by MS in these studies may occur only when it is being secreted in high levels and has not yet had the time to be processed to PA63. As such, any measures with PA83 are not useful in predicting anthrax stages.

Phase-2 of slow progression was the most important feature characterized here, since once it is passed, death followed rapidly, irrespective of time. Phase-2 appeared to correspond with the intermediate-progressive clinical stage where the host battles the organism and the effects of intoxication begin to accumulate but have not yet surpassed host defenses. Of all the toxins, LF was the most stable (least variable) during this period, with only one order of magnitude range, from 4.2 to 41.5 ng/mL, for all 11 animals and all 58 time-points comprising phase-2. Related toxins, PA and LTx, were also less variable (S3 Fig). The level of stability of LF during phase-2 suggests the possibility that levels are somewhat controlled, either by the host or organism (or both), held below the threshold until host defenses are overcome and disease progresses to the rapidly fatal phase-3. Bacteremia and ETx were the most variable during phase 2. Lower levels seen for ETx may be influenced more by the fluctuations in bacteria and dynamics of intoxication during phase-2.

In phase-3, whether fast or slow progression, toxins and bacteremia increase rapidly after passing the phase-2 threshold. So, any toxin levels or bacteremia that are higher than the thresholds suggest there are less than 20 hours of life remaining. The higher the levels, the shorter the interval to death (Fig 5B). In phase-1, lower toxin levels correlate with higher toxin ratios. Conversely, in phase-3 higher toxin levels correlate with lower toxin ratios. This includes all ratios except PA/LF which doesn’t change in a consistent manner through the stages of progression. The threshold for LF/LTx is 2.5-5.2 in early stages but declines to 1.7 at the phase-2 threshold and declines further to 1.2 at terminal. It was even as low as 1.0 (100% LTx) in some animals with high toxemia (Table 5). This indicates that most of the LF is present as LTx at terminal stage.

#### Phase-2 threshold and the point of no return

In 1954, using intradermal *B. anthracis* infection in the guinea pig model, Smith and Keppie described a critical level for bacteremia, a point of no return, above which animals died even after treatment had cleared the organism (11, 63). Animals recovered if treated when bacteremia was 0.7 to 1.5x10^6^ cfu/mL and died when it was greater than the critical level (3x10^6^ cfu/mL). The threshold for bacteremia in cynomolgus macaques at 1.6E+05 cfu/mL was 20-fold lower than that described in guinea pigs. This is not surprising for different models with different infection routes. Smith and Keppie also described the relationship between the bacterial count in the blood and the time-dependent interval to death. We observed a similar relationship in cynomolgus macaques, a correlation of toxins and bacteremia with survival time. We also determined the phase-2 threshold for entry to phase-3. Medical intervention is usually unsuccessful in the late-fulminant stage-3 (33) (equivalent to phase-3), the phase-2 thresholds could represent the potential point of no return for treatment failures.

## Conclusions

The assessment of all major anthrax toxins and bacteremia, along with hematology and pathology allowed a comprehensive evaluation of the natural history of inhalation anthrax in the cynomolgus macaque NHP model. We defined a triphasic kinetics of toxemia which could be aligned with the three clinical stages of anthrax. This alignment for fast and slow survival groups indicated that anthrax progression is toxin-dependent rather than time-dependent. Hematological fitness factors may influence the disease course and will be evaluated in other studies. Lethal factor represented the most significant indicator of anthrax progression with a remarkably invariable phase-2, such that a small increase potentially advances infection to phase-3 and a relatively rapid death. Phase-2 thresholds identified, represent a potential point of no return. Ongoing studies will test the critical thresholds for predicting antimicrobial treatment failures. Future studies with combined antimicrobial and antitoxin treatment may advance our understanding of anthrax in humans and when specialized treatments are most efficacious.

## Methods

### Reagents

Materials and reagents were obtained from Sigma-Aldrich (St Charles, MO) except where indicated. LF, PA83, and PA63 were obtained from List Biological Laboratories (Campbell, CA) and EF was from Dr. Stephen Leppla’ s laboratory NIH/NIAID as described (23). All other reagents, materials and methods for anthrax toxin measurements were described previously (22, 23, 44, 45, 64).

### Safety and quality

The animal exposure study was conducted in 2014 at Battelle Biomedical Research Center (Columbus, OH) in a BSL3 secured laboratory registered with the Centers for Disease Control and Prevention (CDC) for select agents and inspected by the Department of Defense and U.S. Department of Agriculture. All requirements for select agent laboratories and handling of *B. anthracis* were followed. Battelle performed all aspects of the animal study as well as sample analyses for quantitative bacteremia and hematology as described in detail below. Samples sent from Battelle to the CDC for analysis of anthrax toxins were filter sterilized and sterility confirmed prior to shipment. Sample analysis at CDC was performed in a BSL2 laboratory and followed all safety requirements for handling biological specimens. All personnel have full training required for safety and follow all general laboratory, chemical and bloodborne pathogens procedures. Chemical hygiene plans are in place. All sample analyses followed high quality procedures using validated methods and conducted in a CLIA-certified laboratory following strict adherence of CLIA regulations. Battelle follows good laboratory practice guidelines as available in their standard operating procedures and facility practices.

### Animal study

The animal exposure study was conducted at Battelle (Columbus, OH), a Public Health Service (PHS) Animal Welfare Assurance approved facility. The study protocol was approved by the Institutional Animal Care and Use Committee (IACUC) (Approved Protocol #3013) as required by PHS policy following HHS, NIH, Office of Laboratory Animal Welfare principles. Work with animals followed all safety and select agent guidelines described above. All aspects of the animal study protocols were designed to minimize stress in the animals. CDC received study samples from Battelle for toxin analysis and was not involved with any component of the animal protocol. Chief of CDC Animal Care confirmed that the use that Battelle IACUC was sufficient.

Twenty-three *Macaca fascicularis* cynomolgus macaques (12 males, 11 females) from Covance, Inc., Alice, TX, weighing 3.1-4.9 kg prior to spore exposure, were confirmed to be healthy, negative for tuberculosis, seronegative for Simian Immunodeficiency Virus, Simian T-Lymphotropic Virus-1, and Macacine herpesvirus 1 and PCR negative for Simian Retrovirus 1 & 2. Animals were quarantined for 5 weeks in paired housing, then acclimated to single housing in the biosafety level 3 (BSL3) laboratory one week before exposure.

This study was conducted in two challenge groups with the first exposures (Group-1) taking place in November 2013 and the second in February 2014 (Group-2). Group-1 included 10 animals of which 8 (4 males, 4 females) were challenged and 2 (1 male, 1 female) were not challenged (naïve controls). In Group-2, 15 CM (8 males, 7 females), which included the 2 controls from Group-1, were challenged.

Spore challenge exposures in this model were performed as described previously with a target dose of 200 median lethal dose (LD_50_) *B. anthracis* Ames spores via head-only inhalation exposure, and measurement of actual spore dose per animal by plethysmography (37, 42).

Animals were routinely monitored, and clinical signs recorded twice-daily pre-challenge and every 6±1 hours commencing at 12 hours post-challenge. The following clinical signs of anthrax were monitored, anorexia, lethargy, respiratory distress, moribundity, abnormal activity such as recumbency, weak, or unresponsive, as well as seizures or any other abnormal signs. Signs of symptoms recorded included normal (N), hunched posture (HP), no stool (NS), soft stool (SS), diarrhea (DI), vomiting (V), lethargic (L), seizures (SE), labored breathing (LB), wheezing (W), moribund (M), unresponsive (UR), prostrate (P). Animals found to be moribund were humanely euthanized. Time to symptom onset was the difference in time hours and minutes from challenge to the hours and minutes of first specific clinical sign described from the 6-hour monitoring during the post-challenge period. NS was not included as a specific sign since it was also described prior to exposure and was not specific.

Body temperatures for all animals including group-1 controls were monitored via an implantable programmable temperature transponder (IPTT-300 BMDS, Seaford, DE), one each between the shoulder blades and left hip. Temperatures were recorded daily from day -7 through 0 (before exposure) and every 6±1 hours post-challenge. The threshold for a significant increase in body temperature (SIBT) was previously established as the mean pre-challenge (or baseline) temperature plus 2 standard deviations (SD) of the pre-challenge temperature (47).

### Sample collection

Samples collected for Group-1 included whole blood for quantitative bacteremia and plasma for toxins in EDTA tubes at -7 days pre-exposure (-7D) and post-exposure at 18-, 24-, 30-, 36-, 48-, 60-, 72-, 84-, and 96-hours and terminal sample when possible. Plasma sample collection was adjusted for Group-2 animals to include a 42-hour collection time and additional samples at 108-, 120-, 132-, and 144-hours and end of study (day 14). Whole blood for critical blood counts (CBC) (hematology) was collected in EDTA tubes for Group-1 animals at -7D, 18-, 24-, 30-, 36-, 48-, and 72-hours and for Group-2 at -7D, 18-, 24-, 30-, 36-, 48-, 72-, 84-, 96-, 108-, 120-, and 144-hours post-challenge.

### Quantitative bacteremia

The concentration in colony forming units (CFU) of *B. anthracis* vegetative bacteria per mL of whole blood was determined by serial dilution, plating and colony enumeration as described previously (65). Briefly, 100 μL each whole blood sample was diluted in 900 μL phosphate-buffered saline (PBS), then serially diluted 10-fold in PBS up to 10^7^. Then 100 μL of each sample and dilutions were plated in triplicate onto Tryptic Soy Agar, incubated at 37°C for 16-24 h. Mean colony counts from 25-250 cfu/mL, below the quantitative range, were approved for reporting by the Study Director and used here as reported. The quantitative LOD was 100 cfu/mL. Values down to 25 cfu/mL when reported were included. A result of ‘+’ indicated that the mean colony counts for a dilution set (2 of 3 plates) was less than 10. For within group quantification, such as at symptom onset, values of ‘+’ were assigned the qualitative bacteremia LOD of 6 cfu/mL (46) and negative samples were assigned a value at ½ the LOD, 3 cfu/mL.

### Hematology

Hematology, CBC (whole blood) measurements were determined at Battelle with the Advia® 120 (Siemens, Deerfield, IL). Measurements included, white blood cell count and differential leukocyte count (absolute and percentage), hemoglobin, hematocrit, red blood cell count, mean corpuscular volume (MCV), mean corpuscular hemoglobin, mean corpuscular hemoglobin concentration (MCHC), corpuscular hemoglobin concentration mean (CHCM), red cell distribution width (RDW), platelet count, mean platelet volume (MPV), neutrophil:lymphocyte ratio (N/L ratio).

### Mass spectrometry (MS) methods for anthrax toxins

MS methods for anthrax toxins were conducted at CDC. Validated methods for analysis of total-LF and LTx have been described previously (22, 44). Activity of EF monomer and ETx (PA_63_-EF) are congruent and were analyzed for EF activity as described (23). Analysis of total-PA and PA83 for select tryptic peptides in the PA20 and PA63 domains was also performed as described (45). The limits of detection for all the anthrax toxin methods are included in Table 2. Performance characteristics are detailed in the method publications (22, 23, 44, 45).

Anthrax toxin MS methods provide high sensitivity, specificity, and accurate measures of all anthrax toxins (22, 45). Magnetic monoclonal antibody bead capture allows selective immunoprecipitation of toxins, (Step 1), followed by endoproteinase activity of LF/LTx, adenylate cyclase activity of EF/ETx and on-bead tryptic digest for PA (Step 2), and detection of LF/LTx and EF/ETx catalytic products and PA tryptic peptides by isotope-dilution MS (Step 3). The detection of catalytic products of LF, LTx, EF and ETx activity, facilitates signal amplification and enables exquisite analytic sensitivity. LC-MS/MS quantification of specific PA tryptic peptides allows accurate quantification of both total-PA and PA83, and determination of the amount of active PA63. These methods were used to measure toxins in cynomolgus macaques throughout infection. Throughout this report, total LF is referred to as LF and the amount of LF in complex with PA63 as LTx. Similarly, total EF is EF and in complex as ETx. Total PA is described as PA and the individual components as PA83 or PA63.

### Pathology

All cynomolgus macaques, including the survivor to the end of the study, underwent complete necropsy with recording of gross observations. The brain, kidney liver, lung, bronchial, mandibular, and mesenteric lymph nodes, spleen, and gross lesions were collected and preserved in 10% neutral buffered formalin. Preserved tissues were trimmed, paraffin-embedded, sectioned to approximately 5 microns in thickness, affixed to glass slides, and stained with hematoxylin and eosin for brightfield microscopic examination by a board-certified veterinary pathologist. Microscopic findings were graded semi-quantitatively using the following scale: minimal (grade 1) for a minimally detectable lesion, mild (grade 2) for an easily visualized lesion, moderate (grade 3) for a lesion affecting larger areas of the tissue and marked (grade 4) for lesions affecting maximum areas of the tissue. Group summary incidence and average severity calculations were performed for fast and slow progression cohorts. Total Bacteria scores were calculated for each animal by adding the individual microscopic severity scores for bacteria in all examined organs. Total Pathology scores were calculated for each animal by adding the individual microscopic severity scores for all anthrax-related lesions (except for bacteria) in all examined organs as well as the incidence of any gross lesions in the animal.

### Statistical Analysis and Reporting

Toxin levels for total LF (LF), LTx, total EF (EF) and ETx were measured and reported in ng/mL. Total PA (PA) and PA83 were measured in pmol/mL and converted to ng/mL. Calculations of total PA, PA63 and PA83 were determined as follows: In pmol/mL total PA minus PA83 gives PA63 (pmol/mL). PA83 (83 kDa) and PA63 (63 kDa) pmol/mL value multiplied by unit adjusted 83 and 63 ng/pmol, respectively give ng/mL values each which were added to give total PA (PA63+PA83) in ng/mL. Total PA is referred to as PA and any references to PA63 and PA83 are specified.

All calculations of mean, standard deviation, median and interquartile range (IQR), 95 percent confidence intervals, and other analyses where reported were performed using standard equations in Microsoft Excel or JMP®, Version 15, SAS Institute Inc., Cary, NC, 1989-2019. Levels of toxins and bacteremia at specific time points or stages often varied by multiple logs and were skewed (nonparametric). Median and interquartile ranges accurately represented the center of data and range. Mann Whitney Wilcoxon Rank Sum Test (Wilcoxon/Kruskal-Wallis Rank Sum Test in JMP) was used to test the probability that two populations are equal for toxin levels between animals with fast vs slow progression (66). P-value (Prob>|Z|) is reported for the normal approximation test for the two groups (Z test statistic. Alpha level set at 0.05.

For hematological parameters, time post-challenge was limited to 48h because only two animals with fast progression had later measurements, prohibiting comparisons beyond 48h. This demonstrates early changes and differences that may contribute to fast and slow progression. Oneway ANOVA was used to compare differences in means between animals with fast and slow progression at specific time points.

JMP Fit model was used to test the effect of hematology over the time period day -7 pre-challenge, 24-, 36- and 48-hours and for progression (fast vs slow). P-value reported for each term was prob > |t| (t-ratio) and for overall effect tests was prob > F (F-ratio). Alpha level set at 0.05

### Progression determination and kinetics of toxemia

Animals fell into two groups defined by the highest toxin levels and bacteremia reached by 48 hours. Toxemia/bacteremia was higher in 11 animals that died early (fast progression) and were lower in 11 animals that survived longer (slow progression) (n=11) and one animal that survived to the end of the study . In four of 23 animals the initial phase-1 rise proceeded directly to terminal phase-3 (monophasic). The phases were primarily triphasic, consisting of an initial rise (phase-1), plateau/decline (phase-2), and final rise (phase-3). Of all biomarkers measured, LF appeared earliest and was the most consistent (least variable) biomarker during the phase-2 plateau. Therefore, the kinetics of LF was used as the model to estimate the stages of each animal. This estimation is limited by the collection time points. Bacteremia was used to evaluate the kinetics for missing toxin measurements at terminal time points.

Phase-1 was defined by the time of first detection of LF during which it rises to a first peak, the end-of-phase-1, before the plateau or decline. The end-of phase-1 was defined as the time point when biomarker rise slowed or stopped and consequently begins phase-2. In phase-2 the phase-2 plateau continued until the final rapid rise to terminal (phase-3). Phase-3 represents the end of the disease course at (or near) death/euthanasia. Except for bacteremia, terminal toxin levels can only be estimated ‘near’ terminal since in each animal, final samples depended on the symptoms and time of euthanasia. Also, unpredictability and speed of final progression sometimes meant end-stage plasma samples were not available. A terminal whole blood sample for bacteremia was collected even when a plasma sample was not.

Note that some animals were missing a sample collection related to a specific stage. Four animals had no phase-2 (monophasic), of which one animal had no sample for toxemia after 36h (no terminal). Two animals with long survival had a long collection gap between the sample prior to the terminal time and terminal samples. Many were missing terminal plasma for toxins. For phase-3 terminal calculations, only a sample measured ≤12h from the death/euthanasia time point were included.

### Phase-2 and phase-2 threshold

In slow progression, the low variability of LF over the duration of phase-2 (< 1-order of magnitude) allowed characterization of this stage as a whole., Phase-2 was characterized for all toxins, bacteremia, and toxin ratios and included all measurements for all time points from the end-of- phase-1 to the end-of- phase-2 from all 11 animals (n=52, 51 for ETx). The median, upper and lower quartiles, and 95% interval (upper limit) for toxins/bacteremia and 5% interval (lower limit) for toxin ratios were determined in JMP. The phase-2 upper 95% intervals represent potential critical toxin thresholds, which when exceeded signal departure from phase-2 and entry to phase-3. Lower 95% interval applies to toxin ratios.

### Time to threshold and time from threshold to terminal

The time to threshold was calculated from a linear regression for the two time points between which the threshold fell for all animals with available points. Animals with phase-2 levels within the typical standard error (≤10%) of the threshold for LF and followed with lower levels for an extended phase-2, the threshold was determined not to have been exceeded and was calculated at the later time when the final rise continued. Five animals were excluded from the threshold calculations. For one animal with fast progression and one with slow progression the threshold was not exceeded, but later (terminal) samples were missing so the time to threshold could not be calculated for toxins (Fig 3, animal 1, 19). Animals 20 and 22 with slow progression had large time-gaps in sample collection (32 and 58 hours) between the last sub-threshold level sample and final/terminal samples, such that calculation of the time to threshold could not be estimated. The survivor (animal 23) never exceeded the threshold. Two animals (Fig 3, animal 7, 10) were excluded from calculations for PA and ETx because they were below the PA, ETx detection limit before the sample that exceeded the threshold. An n=8 is indicated for the fast group for PA and ETx, versus n=10 for other toxins. Mean time to threshold was calculated for 8-11 animals with fast progression and 8-9 animals with slow progression (exact numbers-n in Fig 5). The time from threshold to terminal was determined by subtracting the calculated time to threshold from the terminal time, estimating how long an animal lived after reaching the threshold (time in phase-3).

## Acknowledgements

This work was supported in part by Health and Human Services (HHS) Agencies: Biomedical Advanced Research and Development Authority (BARDA) and Centers for Disease Control and Prevention (CDC) – Center for Preparedness and Response (CPR). We are grateful to Greg Stark, Maya Sternberg, John Rees, and Susan Kuklenyik for statistical support and discussions.

## Disclaimer

The findings and conclusions in this report are those of the authors and do not necessarily represent the views of the Centers for Disease Control and Prevention. Use of trade names is for identification purposes only and does not imply endorsement by the Centers for Disease Control and Prevention, the Public Health Services or the US Department of Health and Human Services.

The authors declare no competing financial interest.

## Abbreviations

Total lethal factor (LF), total edema factor (EF), total protective antigen (PA), full length 83-KDa PA (PA83), active 63-KDa PA (PA63), lethal toxin (LTx), edema toxin (ETx). Non-human primate (NHP). Large unstained cells (LUC), neutrophil lymphocyte ratio (NL Ratio), white blood cells (WBC), red blood cells (RBC), corpuscular hemoglobin concentration mean (CHCM), mean corpuscular hemoglobin (MCH), mean corpuscular hemoglobin concentration (MCHC), mean platelet volume (MPV), mean corpuscular volume (MCV), red cell distribution width (RDW).

## Supplemental Material

In order of their appearance in text:

**S1 Fig.**
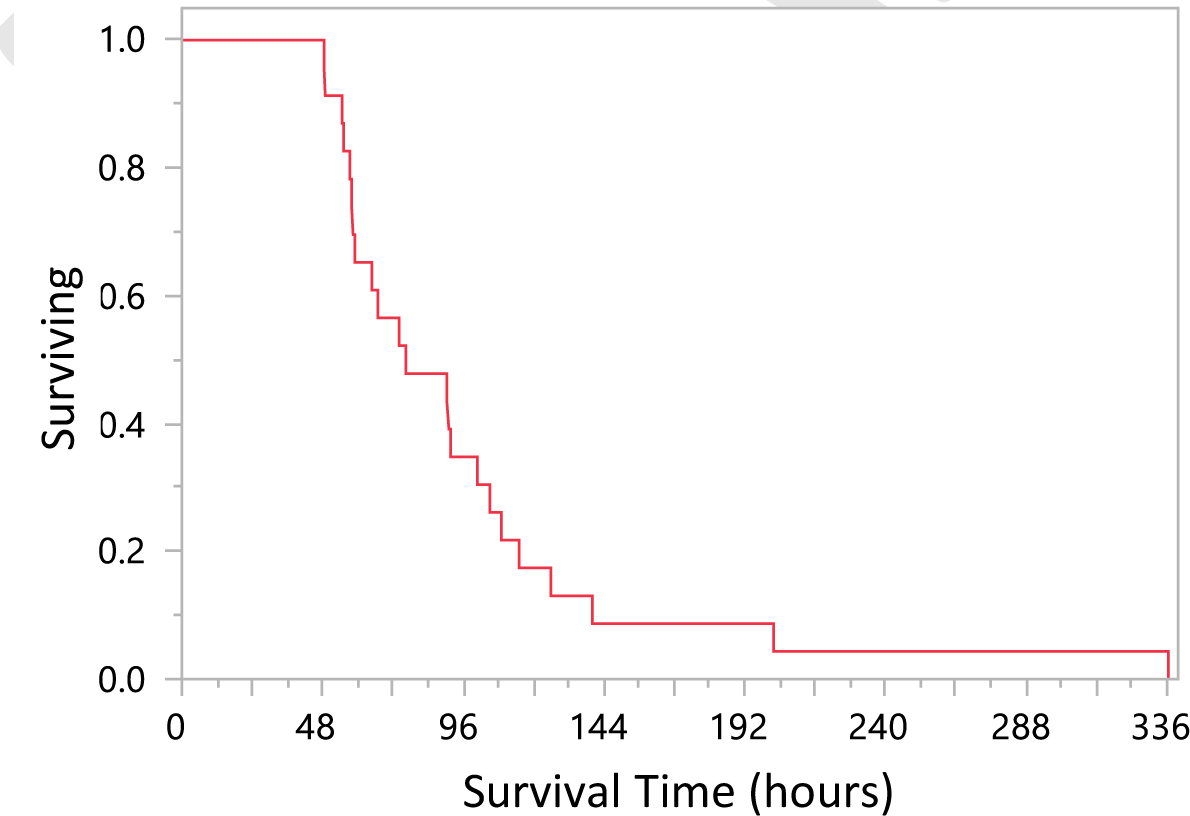
Survival curve for 23 cynomolgus macaques with inhalation anthrax. Proportion surviving vs survival time (hours).

**S2 Fig.**
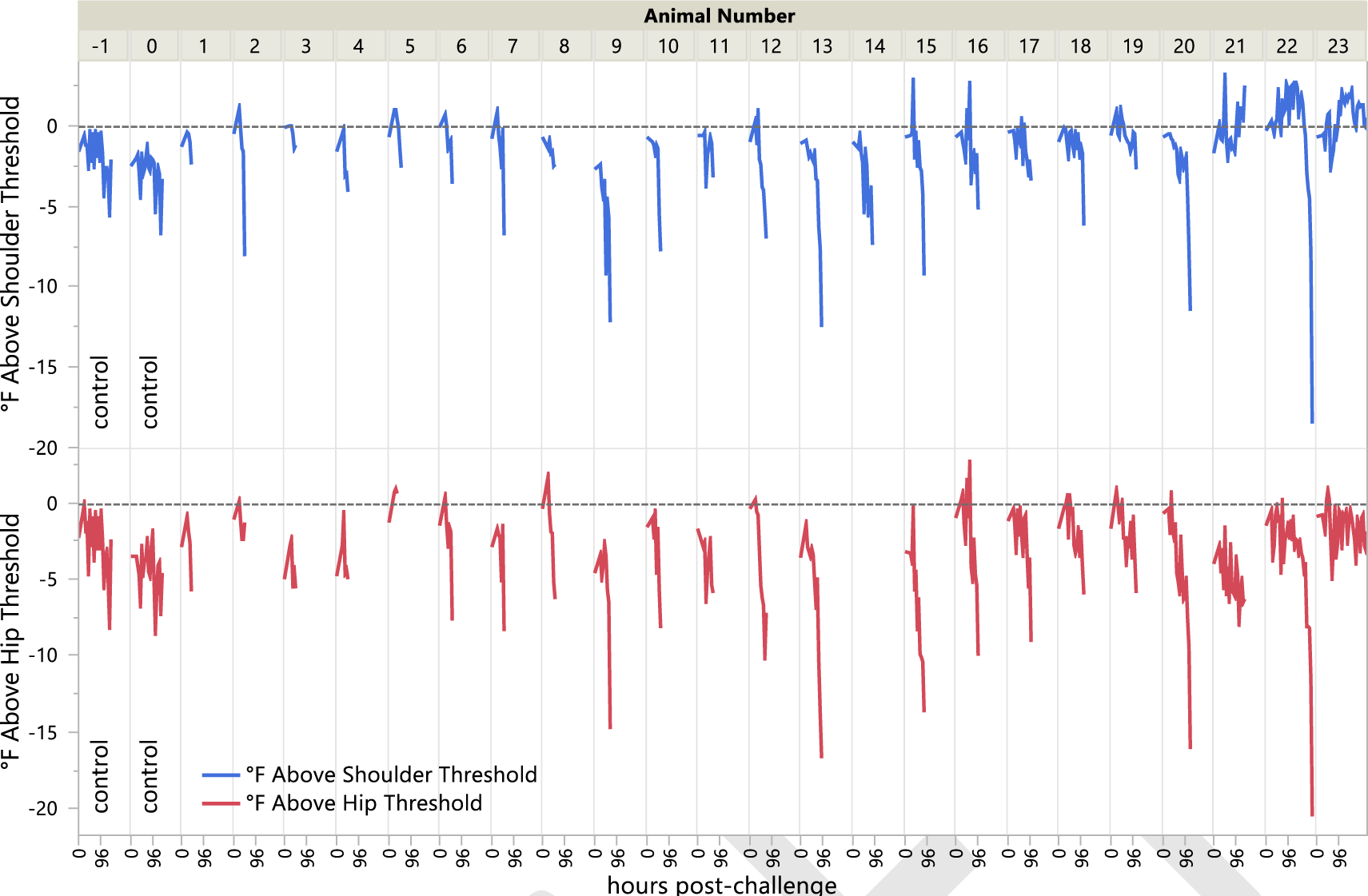
Significant increase in body temperature post-challenge. The threshold for a significant increase in body temperature (SIBT) is established as the mean pre-challenge (or baseline) temperature plus 2 standard deviations (SD) of the mean pre-challenge temperature. Left hip and right shoulder transponders recorded temperatures in °F once daily for seven days. The pre-challenge mean, SD and threshold (mean+2 SD) was determined for both the shoulder and hip transponders for each animal. The deviation from the pre-challenge threshold was calculated by subtracting the normal threshold temperature from the post-challenge temperatures. The solid horizontal line represents the thresholds with zero for the threshold. Deviations from threshold for shoulder and hip graphed for each animal by number according to survival time. Group-1 controls indicated with -1 (C58003) control corresponding to group 2 challenged animal number 5 and 0 (C58170) corresponding to group 2 challenged animal number 11.

S1 File. All anthrax toxin results in cynomolgus macaques with inhalation anthrax.

S2 File. All bacteremia results in cynomolgus macaques with inhalation anthrax.

**S3 Fig.**
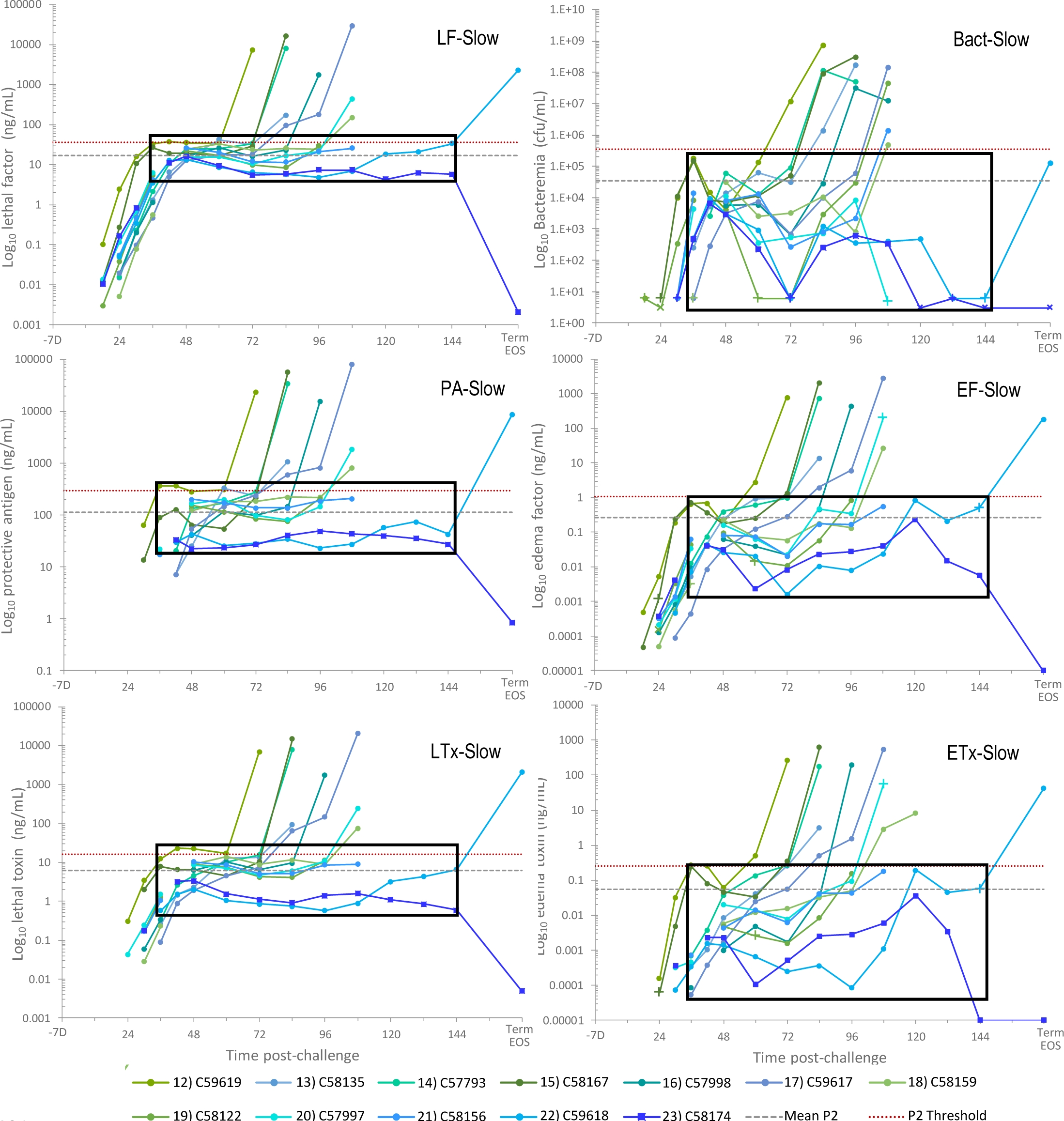
Kinetics of anthrax biomarkers in cynomolgus macaques with slow progression. All toxins (ng/mL) and bacteremia (cfu/mL) over the course of infection in 12 animals with inhalation anthrax and slow progression, of which 11 were triphasic with phase-2 ≥24 hours and one with phase-3 decline and survival. One negative value at the end-of-study (EOS) (animal 23 -C58174) was graphed at ½ the limit of detection for each method as indicated in Table 2. For bacteremia (bact), non-quantifiable results of + was assigned a 6 (+ symbols) and negative at 3 cfu/mL (x symbols). The last time point for C59618 at terminal (Term, 8.4 days) and the survivor which declined to less than the limit of detection at the end-of-study (EOS, 14-days/336 hours) are not to scale. Black boxes represent the range of levels during the phase-2 for each biomarker.

**S1 Table.** End-of-phase-1 toxins, bacteremia, and toxin ratios. Individual results at the end-of-phase- 1(P1) are given for animals with high toxemia, low ratios, and early time-to-death (fast progression) and animals with low toxemia, high ratios, and later time-to-death. Median, upper quartile, and lower quartile, determined in JMP. P-values comparing fast and slow progression obtained by Mann-Whitney-Wilcoxon rank sum (non-parametric) test. Levels for survivor shown separately. *Still in phase-1, excluded. **Less than the limit of detection (<LOD) for PA83 (1.22 ng/mL) given a value at ½ times the LOD for PA83 (0.61 ng/mL) for calculations.

| Animal ID | Time to Death (hours) | End-of-P1 Time Point | Total-PA (ng/mL) | PA83 (ng/mL) | Total LF (ng/mL) | Lethal Toxin (ng/mL) | Total EF (ng/mL) | Edema Toxin (ng/mL) | Bact (cfu/mL) | PA/LF Ratio | LF/LTx Ratio | LF/EF Ratio | EF/ETx Ratio | PA/ETx Ratio | Phase |
| --- | --- | --- | --- | --- | --- | --- | --- | --- | --- | --- | --- | --- | --- | --- | --- |
| 1) C58282* | 48.5 | 48 h NA (36h shown) | 9.11 | <LOD | 2.99 | 0.865 | 0.026 | 0.0045 | 5.3E+02 | 3.0 | 3.5 | 115.0 | 5.8 | 2024 | P1* |
| 2) C59621 | 48.9 | 42 h | 333 | 67.4 | 57.8 | 23.7 | 20.7 | 1.872 | 2.9E+06 | 5.8 | 2.4 | 2.8 | 11.1 | 178 | EOP1-P2 |
| 3) C60763 | 54.7 | 30 h | 326 | 45.2 | 50.9 | 15.3 | 2.6 | 0.576 | 9.4E+05 | 6.4 | 3.3 | 19.6 | 4.5 | 566 | EOP1-P2 |
| 4) C58176 | 55.2 | 30 h | 188 | 25.6 | 200 | 81.7 | 2.68 | 0.635 | 8.3E+06 | 0.9 | 2.4 | 74.6 | 4.2 | 296 | EOP1-P2 |
| 5) C58003 | 57.3 | 42 h | 1143 | 163 | 300 | 169 | 23.7 | 5.64 | 5.9E+06 | 3.8 | 1.8 | 12.7 | 4.2 | 203 | EOP1-P2 |
| 6) C59656 | 57.9 | 48 h | 731 | 182 | 75.2 | 27.7 | 28 | 4.97 | 3.6E+06 | 9.7 | 2.7 | 2.7 | 5.6 | 147 | EOP1-P2 |
| 7) C58214 | 58.4 | 48 h | 509 | 97.6 | 56.2 | 21 | 5.58 | 0.307 | 1.3E+06 | 9.1 | 2.7 | 10.1 | 18.2 | 1658 | EOP1-P2 |
| 8) C60732 | 59 | 36 h | 1189 | 176 | 313 | 165 | 39.2 | 7.1 | 7.4E+06 | 3.8 | 1.9 | 8.0 | 5.5 | 167 | EOP1-P2 |
| 9) C59669 | 64.8 | 48 h | 3035 | 621 | 1038 | 573 | 208 | 38.9 | 3.2E+07 | 2.9 | 1.8 | 5.0 | 5.3 | 78 | EOP1-P2 |
| 10) C57161 | 66.9 | 48 h | 2221 | 516 | 366 | 245 | 53.4 | 15 | 7.2E+06 | 6.1 | 1.5 | 6.9 | 3.6 | 148 | EOP1-P2 |
| 11) C58170 | 74.1 | 48 h | 102 | 12.1 | 38.4 | 5.53 | 2.16 | 0.134 | 3.3E+05 | 2.7 | 6.9 | 17.8 | 16.1 | 761 | EOP1-P2 |
|  | Median |  | 620 | 130 | 138 | 54.7 | 22.2 | 3.4 | 4.8E+06 | 4.8 | 2.4 | 9.0 | 5.4 | 190 |  |
|  | lower quartile |  | 328 | 40.3 | 54.9 | 19.5 | 2.7 | 0.51 | 1.2E+06 | 2.9 | 1.8 | 4.4 | 4.2 | 148 |  |
|  | upper quartile |  | 1447 | 266 | 138 | 188 | 42.8 | 9.1 | 7.6E+06 | 7.1 | 2.9 | 18.2 | 12.3 | 615 |  |
| 12) C59619 | 76.4 | 36 h | 368 | 36.2 | 32.5 | 12.5 | 0.661 | 0.278 | 1.8E+05 | 11.3 | 2.6 | 49.2 | 2.4 | 1324 | EOP1-P2 |
| 13) C58135 | 90.4 | 60 h | 328 | 33.1 | 41.5 | 13 | 0.922 | 0.153 | 6.2E+04 | 7.9 | 3.2 | 45.0 | 6.0 | 2144 | EOP1-P2 |
| 14) C57793 | 91.1 | 48 h | 133 | 8.8 | 23.8 | 4.55 | 0.401 | 0.038 | 6.1E+04 | 5.6 | 5.2 | 59.4 | 10.6 | 3500 | EOP1-P2 |
| 15) C58167 | 91.7 | 36 h | 89.7 | 7.6 | 26.3 | 7.94 | 0.743 | 0.242 | 1.5E+05 | 3.4 | 3.3 | 35.4 | 3.1 | 371 | EOP1-P2 |
| 16) C57998 | 100.8 | 48 h | 41 | <LOD** | 16.2 | 6.32 | 0.064 | 0.001 | 5.6E+03 | 2.5 | 2.6 | 253.1 | 64.0 | 41000 | EOP1-P2 |
| 17) C59617 | 105.1 | 48 h | 54.2 | <LOD** | 12.4 | 1.91 | 0.032 | 0.002 | 3.3E+03 | 4.4 | 6.5 | 387.5 | 16.0 | 27100 | EOP1-P2 |
| 18) C58159 | 109 | 48 h | 121 | 7.89 | 23.8 | 8.76 | 0.231 | 0.0057 | 3.1E+04 | 5.1 | 2.7 | 103.0 | 40.5 | 21228 | EOP1-P2 |
| 19) C58122 | 115.1 | 48 h | 152 | <LOD** | 22 | 9.03 | 0.097 | 0.0048 | 3.6E+03 | 6.9 | 2.4 | 226.8 | 20.2 | 31667 | EOP1-P2 |
| 20) C57997 | 125.9 | 48 h | 165 | 11.8 | 15.6 | 8.62 | 0.166 | 0.020 | 1.1E+04 | 10.6 | 1.8 | 94.0 | 8.3 | 8250 | EOP1-P2 |
| 21) C58156 | 140.1 | 48 h | 200 | <LOD** | 26.1 | 10.3 | 0.081 | 0.0043 | 7.8E+03 | 7.7 | 2.5 | 322.2 | 18.8 | 46512 | EOP1-P2 |
| 22) C59618 | 201.9 | 42 h | 29.6 | <LOD** | 12.4 | 1.53 | 0.046 | 0.0015 | 9.4E+03 | 2.4 | 8.1 | 269.6 | 30.7 | 19733 | EOP1-P2 |
|  | Median |  | 133 | 7.6 | 23.8 | 8.6 | 0.17 | 0.0057 | 1.1E+04 | 5.6 | 2.7 | 103 | 16 | 19733 |  |
|  | lower quartile |  | 54.2 | 0.61 | 15.6 | 4.6 | 0.064 | 0.0020 | 5.6E+03 | 3.4 | 2.5 | 49.2 | 6.0 | 2144 |  |
|  | upper quartile |  | 200 | 11.8 | 26.3 | 10.3 | 0.66 | 0.15 | 6.2E+04 | 7.9 | 5.2 | 270 | 30.7 | 31667 |  |
|  | Fast-v-Slow p-value |  | 0.0035 | 0.0003 | 0.0002 | 0.0011 | 0.0001 | 0.0003 | 0.0001 | 0.5495 | 0.149 | 0.0004 | 0.098 | 0.0004 |  |
| 23) C58174 | Survivor | 42 h | 33.9 | <LOD | 11.1 | 3.25 | 0.040 | 0.0023 | 6.4E+03 | 3.05 | 3.42 | 278 | 17.4 | 14739 | EOP1-P2 |

**S4 Fig.**
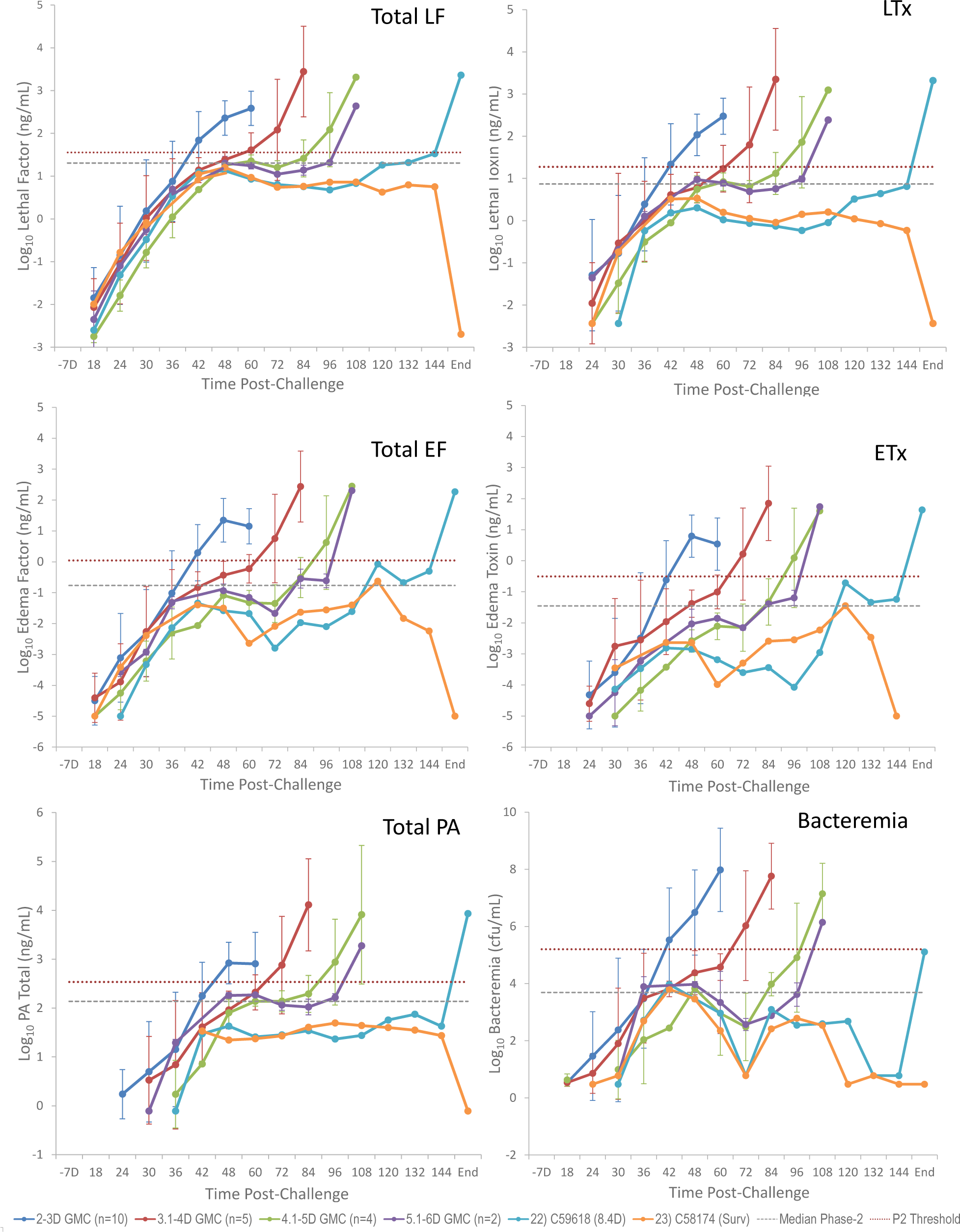
Relationship between anthrax biomarker levels and survival time (days). Mean log_10_ concentrations and standard deviations (error bars) for total LF, LTx, total EF, ETx, total PA, and bacteremia, over the time post-challenge (hours-h) for animals that died/euthanized at 1.9-3.0 days (n=10), 3.1 to 4.0 days (n=5), 4.1 to 5 days (n=4) and 5.1 to 6 days (n=2), and for one animal at 8.4 days (animal 22) and survivor to end-of-study (End) (animal 23). Final time points for animal 22 (C59618) at 8.4 days and for 23 (C58174) at end-of-study (14 days) were included together (End). Gray dashed lines shown for median phase-2 level and phase-2 thresholds (red dotted line).

S3 File. All hematology results in cynomolgus macaques with inhalation anthrax.

